# Exogenous attention generalizes location transfer of perceptual learning in adults with amblyopia

**DOI:** 10.1101/2021.04.27.441700

**Authors:** Mariel Roberts, Marisa Carrasco

## Abstract

Visual perceptual learning (VPL) is a behavioral manifestation of brain neuroplasticity. However, its practical effectiveness is limited because improvements are often specific to the trained conditions and require significant time and effort. It is critical to understand the conditions that promote learning and transfer. Covert endogenous (voluntary) and exogenous (involuntary) spatial attention help overcome VPL location specificity in neurotypical adults, but whether they also do so for people with atypical visual development is unknown. This is the first study to investigate the role of exogenous attention during VPL in adults with amblyopia, an ideal population given their asymmetrically developed, but highly plastic, visual cortex. Here we show that training on a discrimination task leads to improvements in foveal contrast sensitivity, acuity, and stereoacuity. Remarkably, exogenous attention helps generalize learning beyond trained spatial locations. Our findings reveal that attention can enhance the effectiveness of perceptual learning during rehabilitation of visual disorders.

## INTRODUCTION

Visual perceptual learning (VPL) refers to enhancements in perceptual sensitivity or discriminability after extensive practice (reviews: Sagi, 2011; Gilbert and Li, 2012; Dosher and Lu, 2017; Seitz, 2017; Maniglia and Seitz, 2018). There are significant translational implications for training visual expertise and rehabilitation of visual disorders. However, VPL improvements can be modest and are often characterized by specificity (reviews: Sagi, 2011; Gilbert and Li, 2012; Dosher and Lu, 2017; Seitz, 2017; Maniglia and Seitz, 2018). In most cases, performance improvements do not transfer beyond the trained retinal location and stimulus feature, and in some cases, they are specific to the trained eye. Thus, there is significant incentive to design training regimens that promote VPL and its transfer to untrained conditions. Elucidating the specific conditions that promote learning and generalization provides insight into the potential mechanisms and neural substrates underlying plasticity in the (a)typical adult brain. Protocols that require training with a second task, i.e., “piggybacking”, a form of double training (e.g., Hung and Seitz, 2014; Wang et al., 2014) and interleaving tasks (Szpiro et al., 2014), facilitate VPL to untrained locations (e.g., Hung and Seitz, 2014; Szpiro et al., 2014; Wang et al., 2014) and features (e.g., Szpiro et al., 2014). However, these protocols increase training time, which is particularly problematic for populations whose visual systems are already taxed and thus may fatigue more quickly, including adults with amblyopia.

Amblyopia is a neurodevelopmental disorder of the cortex characterized by interocular disparities (Sengpiel, 2014; Levi et al., 2015; Kiorpes and Daw, 2018; Levi, 2020). Although typically considered a foveal disorder (Shooner et al., 2015), its deficits and their neural markers are also present across the visual field (Katz et al., 1984; Hou et al., 2016; Roberts et al., 2016; Pham et al., 2018; Ramesh et al., 2020). Whether due to misalignment of the eyes (strabismic subtype), optical blur (anisometropic subtype), or a reduction in retinal contrast resulting in form deprivation (deprivation subtype), people with amblyopia rely on their stronger eye for most visual tasks while actively suppressing the processing of visual input to their weaker amblyopic eye. However, even their dominant “fellow” eye is impaired relative to those of neurotypical observers (review: Meier and Giaschi, 2017). Despite wearing best optical correction, adults with amblyopia exhibit pronounced impairments in a diversity of tasks, e.g., contrast sensitivity, visual acuity, crowding, position discrimination, spatial interaction, letter recognition, stereopsis, visual search, counting, multiple-object tracking, and oculomotor and hand-eye coordination tasks (reviews: Astle et al., 2011; Levi et al., 2015; Tsirlin et al. 2015; Levi, 2020; Rodán et al., 2020). Thus, the amblyopic brain is a useful model system for investigating neuroplasticity as well as its limits.

Visual training has been shown to reduce many of these deficits, specifically in contrast sensitivity, visual acuity, crowding, position discrimination, spatial interaction, letter recognition, and stereopsis (reviews: Levi and Li, 2009; Astle et al., 2011; Sengpiel, 2014; Levi et al., 2015; Tsirlin et al., 2015; Levi, 2020; Rodán et al., 2020). Though not well understood, VPL has shown to be more generalized in amblyopic than neurotypical adults, often showing transfer to untrained stimulus features (e.g., spatial frequencies, orientations), visual dimensions (e.g., acuity, stereopsis), and to the fellow eye (reviews: Levi and Li, 2009; Astle et al., 2011; Sengpiel, 2014; Levi et al., 2015; Tsirlin et al., 2015; Levi, 2020; Rodán et al., 2020). After training the weaker eye, experimenters can measure differences in VPL improvement and transfer compared to the stronger fellow eye, which provides an informative within-subject comparison condition that is not confounded by individual differences. In addition to their contribution to our understanding of neuro(a)typical brain functioning, findings from VPL studies with amblyopic patients can have translational implications for improving the efficiency of rehabilitation methods for visual disorders.

Covert spatial attention –the selective processing of visual information without accompanying eye movements– improves performance in a variety of detection, discrimination and localization tasks (reviews: Carrasco, 2011; Carrasco, 2014; Carrasco & Barbot, 2015). Recent studies manipulating observers’ covert spatial attention during practice have revealed that selective attention can enable VPL (Szpiro and Carrasco, 2015), and help overcome specificity (Donovan et al., 2015; Donovan and Carrasco, 2018; Donovan et al., 2020) in neurotypical adults. Observers who trained with exogenous (involuntary) attention acquired VPL, but those who trained without focused attention under otherwise identical conditions did not (Szpiro and Carrasco, 2015). Moreover, deploying exogenous (Donovan et al., 2015) or endogenous (voluntary; Donovan and Carrasco, 2018) spatial attention during training facilitates the transfer of improved orientation discrimination to untrained locations. Exogenous attention also facilitates VPL transfer in visual acuity to untrained stimulus locations and features (Donovan et al., 2020). In these studies, to isolate the attentional effect, separate groups of observers were trained with either valid attentional cues, i.e., which focused spatial attention at the target location, or neutral cues, i.e., which evenly spread spatial attention across all spatial locations. All groups were tested with neutral cues before and after training, such that evidence of VPL or transfer was only attributable to attentional allocation during training.

Given that spatial attention cues typically improve behavioral performance in amblyopia (Sharma et al., 2000; Roberts et al., 2016; Pham et al., 2018; Ramesh et al., 2020), and that attention helps overcome VPL location (Donovan et al., 2015; Donovan and Carrasco, 2018; Donovan et al., 2020) and feature (Donovan et al., 2020) specificity in neurotypical adults, here we investigated whether and to what extent exogenous attention facilitates VPL and its transfer in adults with amblyopia. There are alternative hypotheses for this open question. On the one hand, given that people with amblyopia benefit from attention in basic visual tasks as neurotypicals do (Carrasco, 2011; Carrasco, 2014; Carrasco & Barbot, 2015), we could hypothesize that training with attention may also facilitate location and feature transfer. On the other hand, given that some have argued that attention is atypical in amblyopia (as reviewed in Roberts et al., 2016; Pham et al., 2018; Verghese et al., 2019), and differences in the neural correlates of visual and attentional processing in the amblyopic brain –including underactivation in primary visual cortex (review: Joly and Franko, 2014), atypical neural signaling across the visual hierarchy (e.g., Hou et al., 2016; Mortazavi et al., 2020, review: Joly and Franko, 2014), and multiple anomalies in the responsivity and topography of neurons (e.g., Schooner et al., 2015; reviews: Levi, 2013; Joly and Franko, 2014; Kiorpes and Daw, 2018)– we could hypothesize that attention may not interact with VPL in the amblyopic brain in a similar manner as in neurotypicals. Furthermore, given that amblyopic adults already show more generalized improvements than neurotypical adults, it is possible that attention would not be able to further potentiate VPL or its transfer to a measurable degree.

Two groups of adults with strabismic, anisometropic or mixed amblyopia (see **Table S1**) trained using Gabor stimuli set at individuals’ contrast thresholds with their amblyopic eye on a two-alternative forced-choice (2-AFC) orientation discrimination task contingent on observers’ contrast sensitivity (**Figure 1a**; see **Main task trial sequence** under **STAR Methods**). The behavioral effects of both types of attention on this task, as well as their neural correlates, have been well-characterized in neurotypical observers (e.g., Carrasco et al., 2000; Lu and Dosher, 2000; Liu et al, 2005; Pestilli et al., 2009; Herrmann et al., 2010; Lu et al., 2011; Dugué et al., 2016; Dugué et al., 2020; Fernández and Carrasco, 2020; Jigo and Carrasco, 2020), and their behavioral effects have been well-established for special populations (amblyopia(Roberts et al., 2016), ASD(Grubb et al., 2013a; Grubb et al., 2013b), ADHD(Kim et al., 2014; Roberts et al., 2017)).

**Figure 1.**
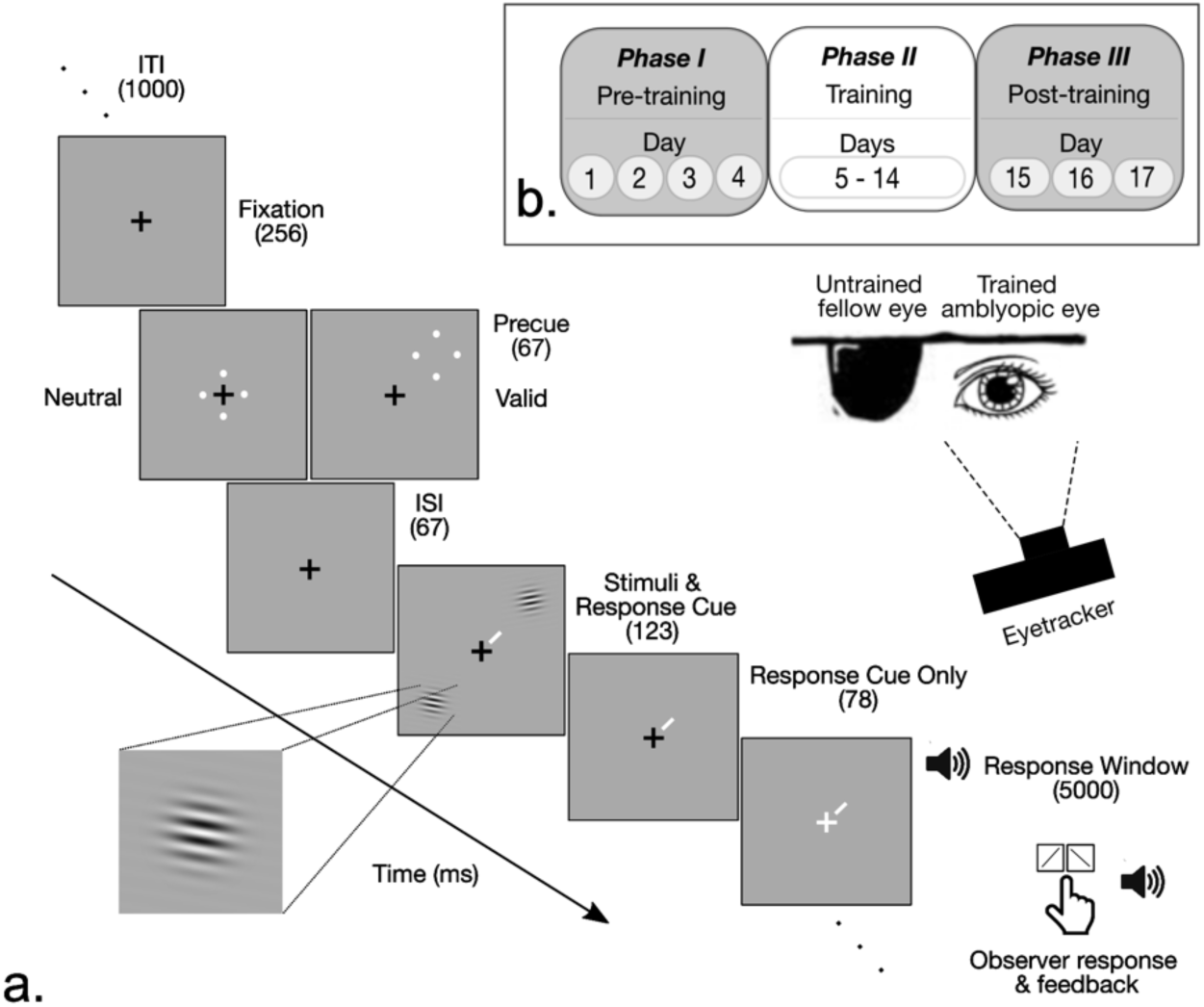
**Experimental protocol. a.** Trial sequence for the main training task. The size of the stimuli has been enlarged for improved visibility. **b.** Overview of study procedure. See also **STAR Methods**.

To assess whether exogenous attention helps overcome location specificity, each group trained (**Figure 1b**) either with peripheral cues (Attention group) or central cues (Neutral group). In the days immediately preceding (Day 4 of the Pre-training phase; **Figure 1b**) and after (Day 15 of the Post-training phase; **Figure 1b**) the training sessions, we measured their performance on the same task in each eye at the trained and untrained diagonal meridians. In addition, to assess the degree of eye, feature and task transfer in both groups, we measured changes in some of the classic visual impairments of amblyopia –broadband foveal contrast sensitivity, visual acuity, crowding at the fovea and binocular stereoacuity– using a curated battery of tasks (see **Supplemental Information** for details).

## RESULTS

### Mean Contrast

We equated performance (∼1.3 d’) between Eyes and Groups before training (**Figure S1**, top row) using an adaptive staircase procedure to estimate monocular contrast thresholds. A 2-way (Eye: amblyopic, fellow X Group: Neutral, Attention) mixed-design ANOVA of mean contrast threshold revealed a significant main effect of Eye (*F*(1,18)=11.7, *p*=.003, η^2^*_G_* =.215), but neither a significant main effect of Group (*F*(1,18)=3.09, *p*=.096, η^2^*_G_* =.091) nor a significant interaction (*F*(1,18)<1). Both the Neutral and Attention groups required about twice the contrast in their weaker amblyopic eye (*M*=35.0±5.16%) than the corresponding fellow eye (*M*=17.0±2.98%) to obtain similar performance. There were no significant correlations between stimulus contrast and normalized improvement –the difference between post-test and pre-test performance divided by the sum of their values (see **STAR Methods**)– at the trained or untrained diagonal for either eye of each group (all *p*>.05).

### Main task performance

We first assessed whether and how exposure to a valid peripheral precue during training differentially affects the magnitude and specificity of learning. We conducted a 4-way mixed ANOVA on d’ for all observers (**Figures S1-S3**, top row) with factors of Session (pre-test, post-test), Eye (amblyopic, fellow), Diagonal (trained, untrained) and Group (Neutral, Attention). There were significant main effects of Session (*F*(1,18)=5.97, *p*=.025, η^2^*_G_* =.058) and Diagonal (*F*(1,18)=5.55, *p*=.030, η^2^*_G_* = .018), and a 3-way interaction among Session, Diagonal, and Group (*F*(1,18)=4.39, *p*=.050, η^2^*_G_* =.006). Neither the main effects of Eye or Group nor all other interactions were significant (all *p*>0.3). The complementary 3-way (Eye X Diagonal X Group) mixed ANOVA of normalized change revealed a significant 2-way interaction between Diagonal and Group (*F*(1,18)=7.00, *p*=.016, η^2^*_G_*=.042); the Neutral group exhibited significantly more improvement at the trained than untrained diagonal (*t*(9)=3.19, *p*=.011, 95% CI=[.044,.260]), whereas the Attention group improved to a similar extent at both diagonals (*t*(9)=-.031, *p*=.976, 95% CI [-.146,.142]). None of the main effects or other interactions were significant (all *p*>.1).

A corresponding analysis of median RT, our secondary variable, showed a significant main effect of Session (*F*(1,18)=28.4, *p*<.001, η^2^*_G_*=.112), but no other significant main effects or interactions (all *p*>.2; **Figures S1-S3**, bottom row). The complementary 3-way (Eye X Diagonal X Group) mixed ANOVA of normalized change indicated a significant Eye x Diagonal interaction (*F*(1,18)=4.88, *p*<.05, η^2^*_G_*=.010), but post-hoc t-tests found no significant differences across all diagonals in both eyes (all *p*>.2). All other main effects and interactions were not significant (all *p*>.1).

Given that the 4-way ANOVA did not reveal a main effect of eye, we collapsed across the amblyopic and fellow eye data for the following analyses. **Figure 2** plots post-test versus pre-test performance (top row: d’, bottom row: RT) collapsed across both eyes of individual observers at the trained (left column) and untrained (middle column) diagonals, as well as the normalized change in performance at one diagonal relative to the other (right column). Training improved performance for most observers at the trained diagonal, but three observers in each group got worse (i.e., exhibited a negative normalized change in d’ at the trained diagonal of the main task). Most critically, all of the observers who exhibited a positive normalized change in d’ at the trained diagonal of the main task exhibited transfer to the untrained diagonal in the Attention group (7/7, 100%) but less than half did so in the Neutral group (3/7, 43%). Further, the normalized change in d’ was more often similar at the trained and untrained diagonals for the Attention group than the Neutral group, as shown by the majority of green shapes clustered near the unity line compared to the majority of blue shapes clustered near the horizontal line in the right column of **Figure 2**.

**Figure 2.**
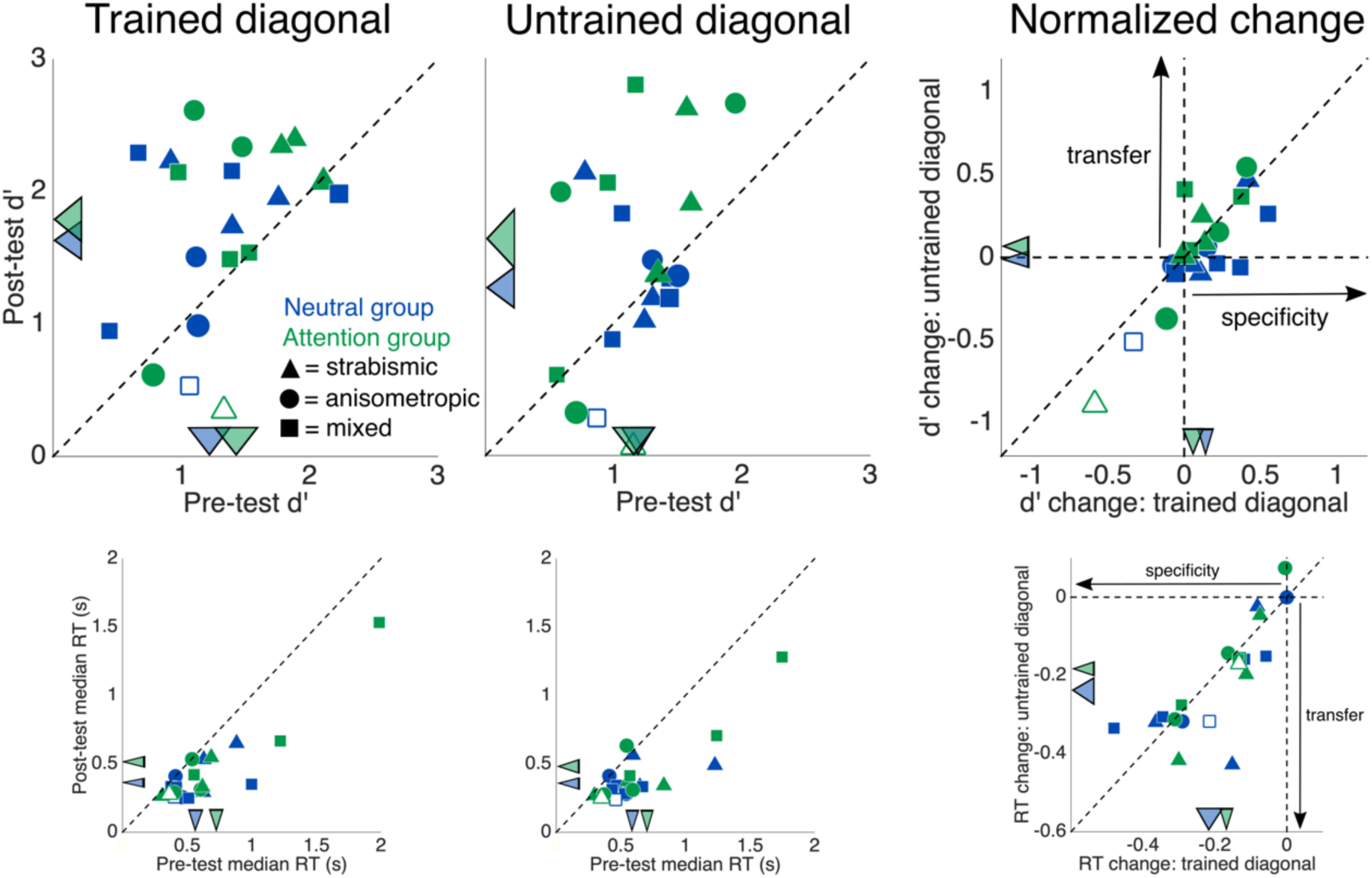
Performance (top row: d’, bottom row: RT) collapsed across both eyes of 20 individual observers on the training task at post-test versus pre-test at the trained diagonal (left column), untrained diagonal (middle column), and the relative normalized change at each diagonal (right column; data to the right of the vertical line indicate improvement at the trained diagonal; data above the horizontal line but below the unity line indicate partial transfer; data along the unity line indicate complete transfer; data above the unity line reveal greater improvement at the untrained compared to the trained diagonal). The colored arrows pointing to the axes indicate group means, and their widths represent ±1 within-groups SEM for each condition (Morey, 2008). Unfilled shapes indicate the one observer per group not included in the statistical analyses for the main training task. See also **Figures S1, S2, and S3**.

RTs were faster for most observers at post-test than pre-test for both the trained diagonal (left column) and untrained diagonal (middle column). Overall, the benefit was similar at both the trained and untrained diagonals for observers in both groups; for the Attention group the data were close to the unity line (right column); for the Neutral group they were more disperse. Thus, we can rule out speed-accuracy tradeoffs for all observers.

### Role of amblyopic subtype on main task performance

We conducted a complementary 4-way mixed ANOVA of observers’ d’ performance on the main task that included one novel between-groups factor, “Subtype” with three levels: strabismic (n=7), mixed (n=8), or anisometropic (n=5). Neither the main effect of subtype (*p*=.172) nor any of its interactions with the other factors of Session, Diagonal, or Group (all *p*>.1) were significant. Thus, our main results did not differ according to observer’s specific subtype of amblyopia, which, among other differences, are associated with varying levels of fixation instability.

### Role of fixation ability on main task performance

To more directly assess the potential effects of observers’ (in)ability to properly fixate on performance in the main task, we conducted another complementary analysis. We first did a median split, categorizing observers into one of two groups according to the number of times their amblyopic eye broke fixation during the post-test at the trained diagonal: “better fixators” were the ten observers who made the fewest fixation breaks and “worse fixators” were the ten observers who made the most fixation breaks. We then conducted a 4-way mixed ANOVA of observers’ amblyopic eye d’ performance on the main task that included a new between-groups factor, “Fixation ability”, with two levels: “better fixators” and “worse fixators”. Neither the main effect of Fixation ability (*p*=.931), nor any of its interactions with the other factors of Session, Diagonal, or Group (all *p*>.1) were significant. Further, the correlation between the number of amblyopic eye fixation breaks and d’ performance at post-test for the trained diagonal was not significant (*p*=.453).

### VPL transfer to untrained locations

To evaluate VPL transfer, logically, an observer should first show learning at the trained conditions. We converted observers’ normalized change in d’ at the trained diagonal (collapsing across the two eyes) into z-scores (relative to each observer’s assigned group mean) and ranked observers in ascending order from those who learned the least to most. We excluded the one observer in the Attention group who could not complete the battery of untrained tasks due to COVID restrictions (open blue circle in all scatter plots) as we could not include this observer in the comparisons between trained and untrained tasks. As this observer also happened to be the one who learned the least, to keep the analysis balanced and include the same number of observers in each group, we excluded the observer in the Neutral group who learned the least. Given that the magnitude of observers’ learning fell along a continuum, ranging from observers who got a bit worse to those who showed a large improvement, we performed a complementary series of analyses to ensure that precluding these two observers who did not improve would not alter our findings. We conducted a complementary analysis in which we systematically precluded the one (**Figure 3**), two (**Figure S4**), or three (**Figure S5**) observers who learned the least from each group. The statistical results were always the same (see **Table S2**). Therefore, in the following two paragraphs we only summarize the detailed results for the analysis in which we dropped the one observer who learned the least in each group (unfilled shapes in all scatterplot figures; both observers’ z-scores < -1.75).

**Figure 3.**
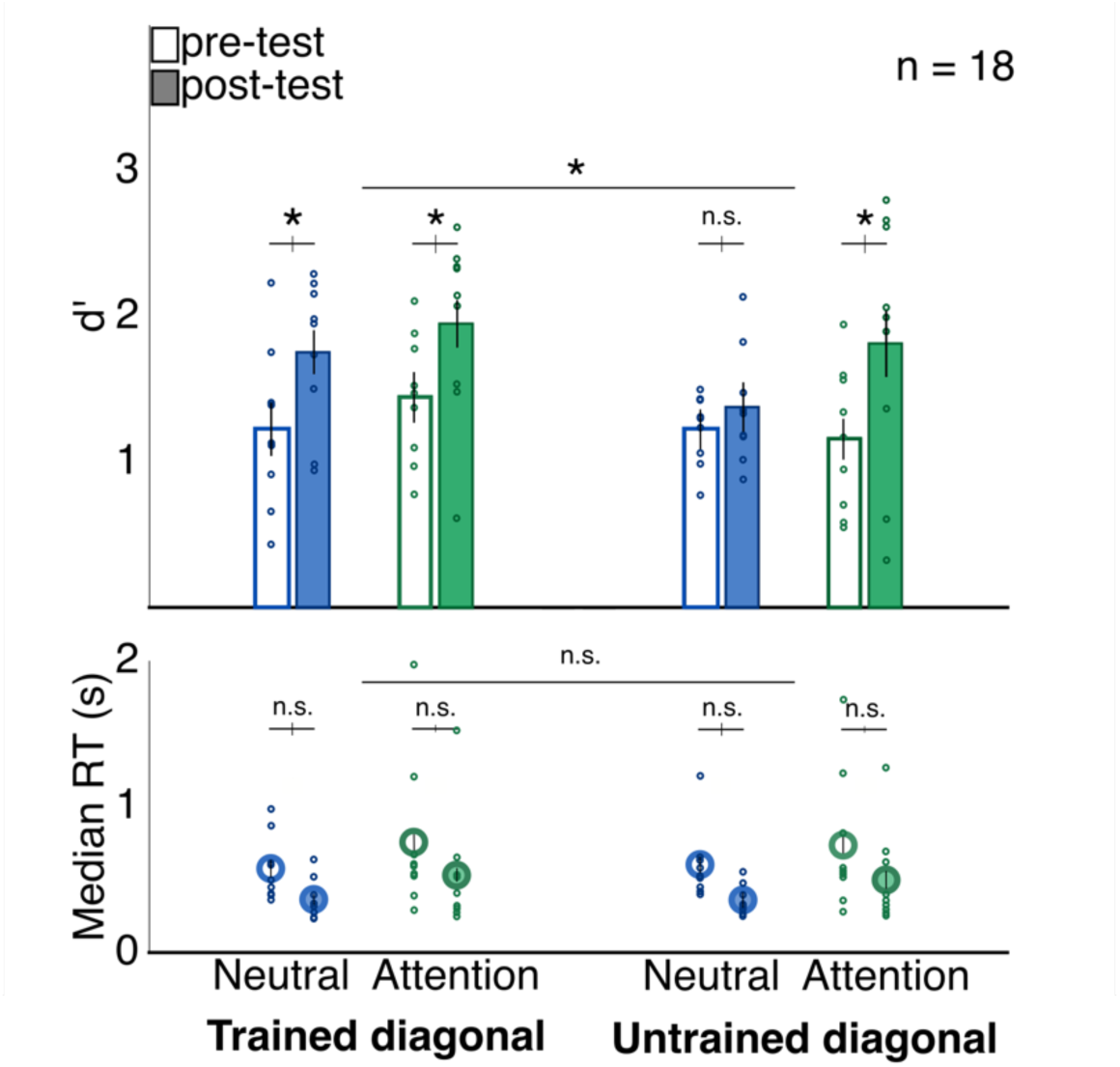
Mean pre-test and post-test performance collapsed across eyes and split by diagonal. Dots indicate the performance of individual observers. Error bars represent ±1 within-subject standard error (Morey, 2008). \**p*<.05. See also **Figures S4**-**S5** and **Table S2**.

To assess whether training with exogenous attention promotes generalization to untrained spatial locations, we conducted a 3-way (Session: pre-test, post-test X Diagonal: trained, untrained X Group: Neutral, Attention) mixed ANOVA of d’ in each group (**Figure 3**, top row). Critically, there was a significant 3-way interaction among Session, Diagonal and Group (*F*(1,16)=4.96, *p*=.041, η^2^*_G_* =.016). 2-way (Session X Diagonal) ANOVAs within each group revealed that whereas the Neutral group showed a significant Session x Diagonal interaction (*F*(1,8)=9.44, *p*=.015, η^2^*_G_* =.049), the Attention group only exhibited a main effect of Session (*F*(1,8)=9.49, *p*=.015, η^2^*_G_* =.190). Consistent with these findings, the complementary 2-way (Diagonal X Group) mixed ANOVA of normalized change revealed that only the 2-way interaction between Diagonal (trained, untrained) and Group (Neutral, Attention) was significant (*F*(1,16)=5.23, *p*=.036, η^2^*_G_* =.048).

Corresponding analyses of the secondary variable of median RT revealed no significant main effects or interactions (all *p*>.1; **Figure 3**, bottom row). The 2-way (Diagonal X Group) mixed ANOVA of normalized change in RT revealed no significant main effects or interaction (all *p*>.3).

### VPL transfer to untrained tasks

Several VPL studies with amblyopic adults have reported generalized improvements beyond the trained task for some untrained features (e.g., spatial frequencies and orientations) and visual dimensions (reviews: Levi and Li, 2009; Sengpiel, 2014; Tsirlin et al., 2015; Levi, 2020; Rodán et al., 2020). Thus, we also assessed whether, after training on a contrast sensitivity-dependent orientation discrimination task in the periphery, observers would show improvement in a curated battery of visual tests assessing broadband and letter foveal contrast sensitivity, crowding and stereoacuity.

**Table S3** provides the groups means at pre-test, post-test and in terms of normalized change for all tasks. **Figure 4** provides a summary of the results of a series of 3-way (Eye: amblyopic, fellow X Group: Neutral, Attention X Session: pre-test, post-test) mixed ANOVAs for all untrained tasks, which were carefully chosen to assess some of the classic deficits in amblyopia. In all tasks, the amblyopic eye performed worse than the fellow eye (i.e., significant main effect of Eye), verifying that observers were amblyopic and that there was a significant disparity between their eyes on a variety of visual dimensions. The main effect of Group was never significant, showing that both groups were composed of individuals with similarly severe amblyopia, and thus had equal opportunity to improve with training. Observers significantly improved at post-test for all tasks except for the one that measured the critical spacing of crowding. There was a significant 3-way interaction among Eye, Group and Session for the ‘area under the log qCSF’ (AULCSF), a measure of foveal broadband contrast sensitivity, and a significant 2-way interaction of Eye and Group for the complementary measure of normalized change in the AULCSF (**Figure S8**); the Neutral group only showed modest improvement in their amblyopic eye, whereas the Attention group showed similar but modest improvements in both their amblyopic and fellow eyes. The amblyopic eye improved more than the fellow eye for the Pelli-Robson contrast sensitivity task, a coarser measure of foveal letter contrast sensitivity, as shown by a significant 2-way interaction of Session x Eye and main effect of Eye for normalized change in Pelli-Robson values (**Figure S9**).

**Figure 4a.**
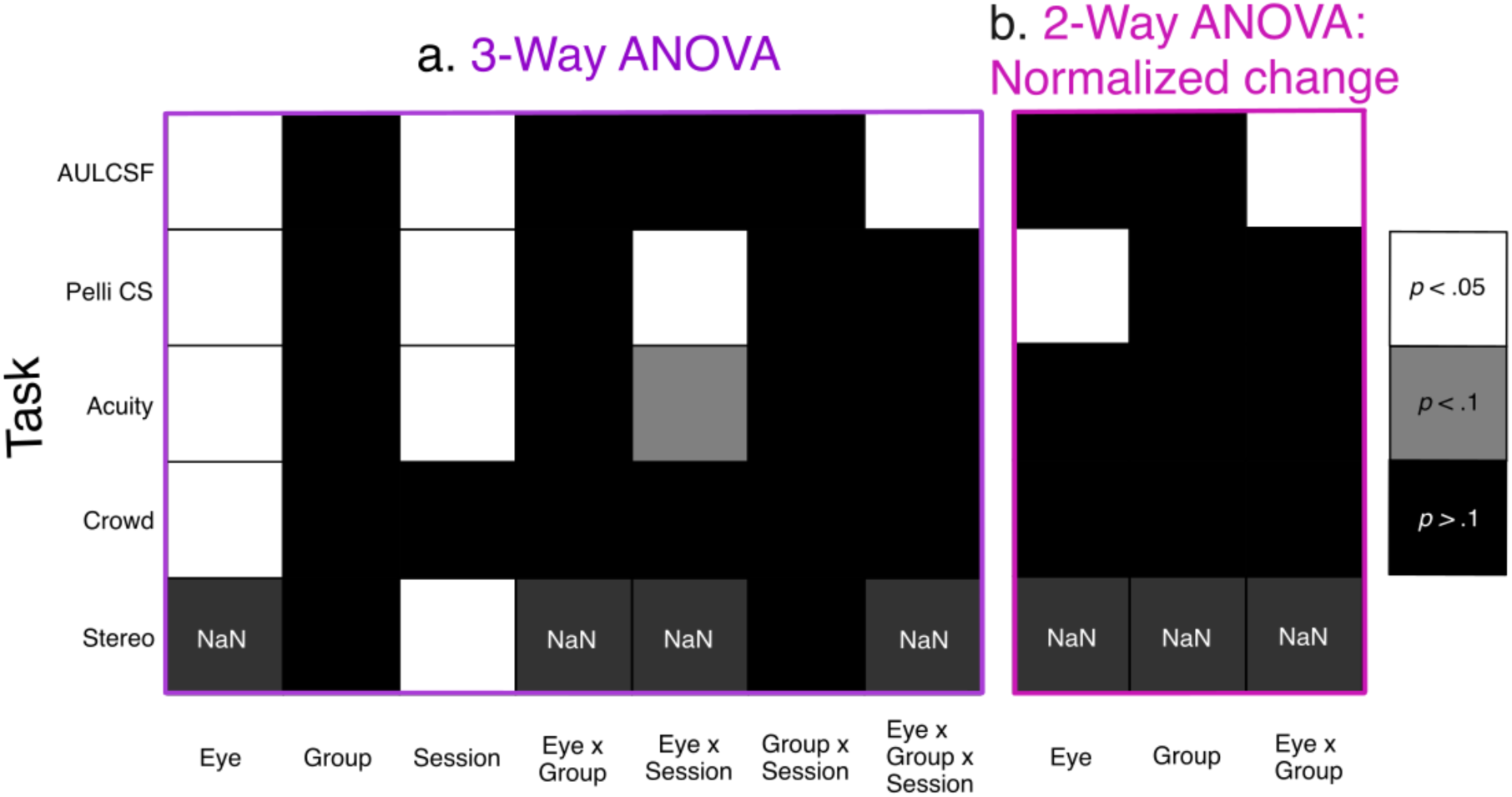
A graphical representation of the results of a series of 3-Way ANOVAs for all untrained visual tasks (left table) and **b.** 2-WAY ANOVAs for their corresponding normalized change values (right table). The shading of each square represents the statistical significance of the main effect or interaction in each of the respective tasks. See also **Table S3, Figures S8-S12**.

### qCSF

To assess potential changes in observers’ broadband contrast sensitivity at the fovea, we had them perform a 4-AFC orientation discrimination task that incorporated the quick Contrast Sensitivity Function (qCSF) procedure on days 2 and 16 (before and after training) (see **Figure S6**). The qCSF is a Bayesian adaptive procedure that uses a trial-by-trial information gain strategy to estimate and detect changes in the underlying contrast sensitivity functions of neurotypical (e.g., Hou et al., 2016) and special populations, including adults with amblyopia (Hou et al., 2010; Jia et al., 2018). Each observer’s contrast sensitivity function was modeled as a truncated log-parabola with four parameters: peak sensitivity, peak spatial frequency, bandwidth, and truncation (see **Table S4** for group means).

Each model parameter was treated as a separate dependent variable in a series of 3-way (Eye: amblyopic, fellow X Group: Neutral, Attention X Session: pre-test, post-test) mixed ANOVAs (**Figure S7a**). The main effect of Eye was always significant (amblyopic eye<fellow eye; all *p*<.015). There was a significant 2-way interaction between Group and Eye for the truncation parameter (*F*(1,17)=5.24, *p*=.035, η^2^*_G_*=.028) because the Neutral group showed a significant difference between the two eyes (*t*(9)=-4.81, *p*<.001, 95% CI [-.187,-.067]), but the Attention group did not (*t*(8)=-.893, *p*=.398, 95 CI% [-.095,.042]). There was also a significant 2-way interaction between Session and Eye for peak sensitivity (*F*(1,17)=8.57, *p*=.009, η^2^*_G_* = .010); whereas there was a significant change in peak sensitivity from pre-test to post-test in the amblyopic eye (*t*(18)=- 2.81, *p*=.012, 95 CI% [-.132,-.019]), there was not in the fellow (*t*(18)=-.775, *p*=.449, 95 CI% [-.074,.034]). All other 2- and 3-way interactions were not significant (all *p*>.3). Complementary 2- way (Eye: amblyopic, fellow X Group: Neutral, Attention) ANOVAs of the normalized change (**Figure S7b**) revealed a significant main effect of Eye for peak sensitivity (*F*(1,17)=7.68, *p*=.013, η^2^*_G_* =.057; amblyopic eye>fellow eye) and peak spatial frequency (*F*(1,17)=7.79, *p*=.013, η^2^*_G_*=.171; fellow eye>amblyopic eye), but no main effect of Group or 2-way interactions (all *p*>.2).

A Pearson’s correlation matrix analysis revealed that, aside from the truncation parameter that did not correlate with any of the other parameters (all *p*>.05) and a nonsignificant correlation between peak gain and bandwidth (*p*=.124), all of the qCSF parameters significantly covaried with each other (all *p*<.05) and with a single value that effectively captures simultaneous changes in all model parameters, the ‘area under the log qCSF’ (AULCSF; all correlations *p*<.001). The AULCSF is a reliable and comprehensive index of a neurotypical (e.g., Hou et al., 2016) or amblyopic (Hou et al., 2010; Jia et al., 2018; Gu et al., 2020) observer’s ‘window of visibility’ as it estimates contrast sensitivity across the wide range of spatial frequencies that compose most visual scenes.

A 3-way (Eye: amblyopic, fellow X Group: Neutral, Attention X Session: pre-test, post-test) mixed ANOVA (**Figure 4a**) of observers’ individual AULCSF values (**Figure S8**) revealed significant main effects of Eye (*F*(1,17)=12.3, *p*=.003, η^2^*_G_*=.203; amblyopic eye *M*=1.23±.052, fellow eye *M*=1.50±.051), Session (*F*(1,17)=4.58, *p*=.047, η^2^*_G_*=.037; pre-test *M*=1.32±.017, post-test *M*=1.42±.045), and a 3-way interaction of Eye, Group, and Session (*F*(1,17)=5.84, *p*=.027, η^2^*_G_* =.007). A 2-way ANOVA within the Neutral group revealed a significant Eye X Session interaction (*F*(1,9)=9.51, *p*=.013, η^2^*_G_*=.010), with greater pre-test to post-test improvement in the amblyopic eye (*t*(9)=2.16, *p*=.060, 95% CI [-.006,.234]) than fellow eye (*t*(9)=.136, *p*=.895, 95% CI [-.094,.106]). A 2-way ANOVA within the Attention group only revealed a main effect of Eye (*F*(1,9)=6.66, *p*=.030, η^2^*_G_*=.229); the main effects of Session and interaction were not significant (both *p*>.1). The complementary 2-way (Eye: amblyopic, fellow X Group: Neutral, Attention) mixed ANOVA of normalized change in the AULCSF (**Figure 4b**) only revealed a significant 2-way interaction of Eye and Group (*F*(1,17)=4.67, *p*=.045, η^2^*_G_*=.045); the amblyopic eye (*M*=.068±.028) improved more than the fellow eye (*M*=-.002±.028) in the Neutral group (*t*(9)=2.51, *p*=.033, 95% CI [.007,.133]), but the improvement was similar in the amblyopic eye (*M*=.058±.037) and fellow eye (*M*=.068±.037) in the Attention group (*t*(8)=-.263, *p*=.799, 95% CI [-.095,.075]).

### Pelli-Robson contrast sensitivity

We measured Pelli-Robson scores from an eye chart as a complementary, but relatively coarse, measure of observers’ foveal broadband contrast sensitivity for letters that are composed of several different spatial frequencies (**Figure S9** for individual scatter plots). A 3-way (Eye: amblyopic, fellow X Group: Neutral, Attention X Session: pre-test, post-test) mixed ANOVA (**Figure 4a**) of observers’ Pelli-Robson contrast sensitivity scores for letters (**Figure S9**) revealed significant main effects of Eye (*F*(1,17)=4.66, *p*=.045, η^2^*_G_*=.078; amblyopic eye *M*=1.77±.039 log contrast threshold, fellow eye *M*=1.89±.035 log contrast threshold), Session (*F*(1,17)=7.92, *p*=.012, η^2^*_G_*=.050; pre-test *M*=1.79±.037 log contrast threshold, post-test *M*=1.87±.027 log contrast threshold), and a 2-way interaction of Eye and Session (*F*(1,17)=4.51, *p*=.049, η^2^*_G_*=.028), as there was a significant change between sessions in the amblyopic eye (*t*(18)=-3.63, *p*=.002, 95% CI [-.237,-.063], but not the fellow eye (*t*(18)=-.383, *p*=.706, 95% CI [-.102,.071]). The main effect of Group, all other 2-way interactions, and the 3-way interaction were not significant (all *p*>.1). The 2-way (Eye: amblyopic, fellow X Group: Neutral, Attention) mixed ANOVA of the normalized change (**Figure 4b**) revealed only a significant main effect of Eye (*F*(1,17)=6.27, *p*=.023, η^2^*_G_*=.148; amblyopic eye *M* =.045±.012, fellow eye *M*=.004±.011); all other main effects and interactions were not significant (all *p*>.1).

### logMar visual acuity

We also investigated whether training on a contrast sensitivity task in the periphery would translate to amblyopic and/or fellow eye improvements in logMar acuity at the fovea, the visual dimension traditionally used to diagnose amblyopia (for group mean values at pre-test, post-test and in terms of normalized change, see **Table S3**). A 3-way (Session: pre-test, post-test X Eye: amblyopic, fellow X Group: Neutral, Attention) mixed ANOVA of all observers’ logMar visual acuity (**Figure S10** for individual scatter plots; higher values are worse) revealed highly significant main effects of Session (*F*(1,17)=21.5, *p*<.001, η^2^*_G_*=.10; pre-test *M*=.271±.05, post-test *M*=.147±.04), Eye (*F*(1,17)=42.2, *p*<.001, η^2^*_G_*=.57; amblyopic eye *M*=.425±.04, fellow eye *M*=-.007±.02), and a marginal 2-way interaction of Session and Eye (*F*(1, 17)=4.25, *p*=.055, η^2^*_G_*=.03), indicating that training on a contrast sensitivity task also improved acuity (**Figure 4a**). None of the main effects or interactions in the 2-way (Eye X Group) mixed ANOVA of observers’ line difference change (pre-test – post-test) was significant (all *p*>.05), indicating that there was similar transfer of improved acuity to the untrained fellow eye in both groups (**Figure 4b**).

We also assessed whether and to what extent training reduced the interocular difference in visual acuity (also known as amblyopic depth), a common metric for quantifying amblyopia severity. We calculated the proportion of amblyopic depth corrected in each observer as the difference in amblyopic depth at post-test compared to pre-test, divided by the amblyopic depth at pre-test. One sample t-tests revealed that visual training corrected for a significant proportion of amblyopic depth and ameliorated amblyopic severity in both groups (Neutral: *M*=.508±.19; *t*(9)=2.70, *p*=.024, 95% CI [.08,.93]; Attention: *M*=.519±.21; *t*(8)=2.48, *p*=.038, 95% CI [.04,1.0]).

### Stereoacuity thresholds

Stereoacuity thresholds, or the distance in arcseconds required to reliably distinguish two objects as being at different planes of depth, are indicative of an observer’s 3D vision. Behaviorally, adults with amblyopia are often stereoblind, and suffer dramatic problems with depth during motor tasks from an early age (reviews: Levi et al., 2015; Levi, 2020). Given that stereoacuity is a binocular measure of depth perception, Eye was not included as a factor in a 2- way mixed ANOVA of stereoacuity thresholds (**Figure 4a**; **Figure S11** for individual scatter plots; higher thresholds are worse). There was a significant main effect of Session (*F*(1, 17)=5.60, *p*=.030, η^2^*_G_* =.050; pre-test *M*=826±98.5 arcsec, post-test *M*=658±107 arcsec), but the main effect of Group and 2-way interaction were not significant (both *p*>.5). A one-way ANOVA of the normalized change in stereoacuity revealed that the between-subjects factor of Group was not significant (*p>.4;* **Figure 4b**).

### Critical spacing of crowding

We measured observers’ critical spacing (degrees of visual angle required to reliably and accurately discriminate a central number from two flanking distractor numbers to the left and right) as a proxy for observers’ deficits in visual crowding. A 3-way (Eye: amblyopic, fellow X Group: Neutral, Attention X Session: pre-test, post-test) mixed ANOVA of observers’ critical spacing (**Figure 4a**; **Figure S12** for individual scatter plots; higher values are worse) only revealed a significant main effect of Eye (*F*(1,17)=11.5, *p*=.003, η^2^*_G_*=.245; amblyopic eye *M*=.374±.076°, fellow eye *M*=.112±.007°). In contrast to all other tasks, there was not a significant main effect of Session (*F*(1, 17)=.770, *p*=.392, η^2^*_G_*=.002; pre-test *M*=.233±.032°, post-test *M*=.253±.045°). All other main effects and interactions were not significant (all *p*>.3). Consistently, none of the main effects or interactions in the 2-way mixed ANOVA of normalized change were significant (all *p*>.7; **Figure 4b**).

### Evidence of generalized learning within the same observers: Positive correlations in magnitude of normalized improvement across certain tasks

To assess whether and how performance changes in one task predict changes in another within the same observer, we calculated a Pearson’s correlation matrix. **Figure S13** is a graphical representation of the significance of correlations of normalized change across all tasks after controlling for multiple comparisons for the amblyopic eye (bottom left triangle) and fellow eye (upper right triangle), combining across both groups. All significant correlations showed a positive relation, i.e., improvement in one was correlated with improvement in another. Improvements in the amblyopic eye were significantly correlated across several tasks, with no significant correlations for the fellow eye. We do not interpret this to mean that normalized changes were not correlated across tasks for the fellow eye. Rather, we think the results are at least partially due to a floor/ceiling effect; as the fellow eye was already relatively high performing compared to the amblyopic eye and had a smaller range for improvement, the normalized change values would be artificially compressed, subsequently diminishing our likelihood of detecting a significant correlation.

Despite the relative variability in how much observers improved overall, there was significant within-subjects consistency across tasks. Notably, improvement at the trained diagonal on the main task was significantly correlated with stereoacuity improvements. Furthermore, greater improvements with the amblyopic eye at the untrained diagonal predicted significantly greater improvements in stereoacuity and crowding. Improvements in amblyopic eye acuity were significantly correlated with improvements in crowding, which in turn were significantly correlated with improvements in stereoacuity. Note, however, that at the group level, mean changes in the critical spacing of crowding did not reach statistical significance. Thus, in general the same observers showed a similar degree of improvement across tasks, evidence for widespread improvement not specific to certain locations, tasks or visual dimensions.

### Amblyopic eye Pelli-Robson contrast sensitivity at pre-test correlates with magnitude of improvement

We explored whether pre-test performance for either eye in any of the tasks correlated with normalized change in the same task or others. After correcting for multiple comparisons using the BH adjustment, there was only a significant correlation between amblyopic eye Pelli-Robson contrast sensitivity for letters at pre-test and normalized change in this task (*p*<.011). Specifically, consistent with previous studies (reviews: Levi and Li, 2009; Astle et al., 2011), observers who had worse Pelli-Robson contrast sensitivity to start improved more.

## DISCUSSION

Two groups of adults with a similar severity of amblyopia trained during approximately ten 1-hour sessions on an orientation discrimination task. Their performance was well equated before training by individually adjusting the stimulus contrast presented to each eye for each observer. The pre-test and post-test were identical for both groups, and during the training days the only difference was that one group trained with valid peripheral exogenous precues (Attention group), whereas the other trained with uninformative neutral precues (Neutral group).

Our training protocol successfully induced VPL. After training the amblyopic eye, both eyes of most observers in both groups exhibited similar performance improvements –higher d’ and faster RT– at the trained diagonal. Training with attention did not potentiate the overall improvement at the trained diagonal, consistent with (Donovan et al., 2015; Donovan and Carrasco, 2018; Donovan et al., 2020), but importantly, training with attention generalized VPL by facilitating location transfer. Our study contributes to a rich VPL literature in both neurotypical humans (reviews: Sagi, 2011; Gibert and Li, 2012; Dosher and Lu, 2017; Seitz, 2017; Maniglia & Seitz, 2018) and those with amblyopia (reviews: Levi and Li, 2009; Astle et al., 2011; Levi et al., 2015; Tsirlin et al., 2015; Rodán et al., 2020; Levi, 2020) demonstrating impressive neuroplasticity in the adult brain.

There is evidence that covert spatial attention remains functionally intact in amblyopia (Roberts et al., 2016; Pham et al., 2018; Ramesh et al., 2020), even though the temporal dynamics of visuospatial processing may be altered (e.g., Verghese et al., 2019; Mortazavi et al., 2020). The primary goal of this study was to assess whether exogenous attention modulates perceptual learning in people with amblyopia, specifically whether it facilitates transfer of discriminability improvements to untrained spatial locations. Our experimental protocol enabled us to evaluate changes in d’ rather than just RT: Observers were instructed to be as accurate rather than as fast as possible and were forced to wait before responding to ensure all observers had adequate time to accrue asymptotic levels of sensory information. Importantly, there were no speed-accuracy tradeoffs –observers were fastest during conditions in which they were most accurate.

There was complete interocular transfer of VPL; both eyes exhibited similar magnitude VPL in both groups of observers. Despite being relatively stronger than the amblyopic eye, the dominant, untrained fellow eyes of adults with amblyopia possess significant room for improvement; they underperform in a variety of visual tasks relative to typically developed eyes (Meier and Giaschi, 2017), which exhibit VPL themselves (reviews: Sagi, 2011; Gibert and Li, 2012; Dosher and Lu, 2017; Seitz, 2017; Maniglia & Seitz, 2018). Some monocular training studies have found eye-specific VPL (review: Sagi, 2011), but others in neurotypical adults (Schoups and Orban, 1996) and patients with visual disorders (Sabesan et al., 2017) have found interocular transfer. Overall, our study strengthens the claim that VPL is not specific to the trained amblyopic eye and leads to significant improvements in the untrained fellow eye (reviews: Astle et al., 2011; Tsirlin et al., 2015; Levi and Li, 2009).

### Exogenous attention helps overcome location specificity in adults with amblyopia

This is the first study to investigate the role of exogenous attention during VPL in adults with amblyopia, who have atypically developed, but highly plastic, visual cortex. Despite similar improvements at the trained diagonal in both groups, the Neutral group exhibited location specificity –no significant learning at the untrained diagonal in either eye. In contrast, the Attention group showed VPL location transfer –similar d’ improvement in both eyes at the trained and untrained diagonals. This novel finding that spatial attention helps overcome location specificity in adults with amblyopia parallels the few studies that have isolated the role of spatial attention during VPL and location transfer in neurotypical adults (Donovan et al., 2015; Donovan and Carrasco, 2018; Donovan et al., 2020).

Both groups were only presented with neutral cues during the pre-test and post-test sessions. The only difference was the attention precue during training –either neutral or peripheral, transiently shifting their attention to the target location. Given that the training locations were known in advance, observers could have allocated their endogenous attention to both potential target locations to improve performance. Any such benefits would have been similar in both groups. Moreover, the 134-ms delay between the peripheral precue and stimuli presentation was too short for endogenous attention to be selectively deployed to the target location (which takes ∼300 ms; Müller and Rabbitt, 1989; Nakayama and Mackeben, 1989; Liu et al., 2007; Giordano et al., 2009). Thus, the fact that both groups show similar performance at the trained diagonal but differential improvement at the untrained can only be attributed to differences in exogenous attentional allocation *during training.* These findings are encouraging, as they demonstrate that neuroplasticity in the adult amblyopic brain may extend further than previously known.

### Transfer to untrained tasks

Consistent with other perceptual learning training studies with amblyopic adults, we found that most observers in both groups exhibited concomitant improvement in complementary untrained measures of the trained visual dimension of contrast sensitivity. They improved in the untrained visual dimensions of logMar visual acuity and stereoacuity, but not in the critical spacing of crowding. In sum, our monocular training protocol exhibited a remarkable degree of generalization of VPL to untrained locations, tasks, visual dimensions, and to the fellow eye. The behavioral and neural markers of amblyopia are present across the visual field, including the parafovea (e.g., Katz et al., 1984; Hou et al., 2016; Roberts et al., 2016; Pham et al., 2018; Ramesh et al., 2020), where we placed our training stimuli as that is where covert attention has typically been investigated (reviews: Carrasco, 2011; 2014; Carrasco & Barbot, 2014). It is encouraging that the VPL benefits extended beyond the parafovea, into their central foveal vision, where the amblyopic deficit is worst (Shooner et al., 2015).

#### Complementary measures of broadband contrast sensitivity improvements at the fovea

Contrast sensitivity improvements measured by the qCSF were largely driven by changes in the peak gain (sensitivity) of the contrast sensitivity function, rather than the spatial frequency corresponding to the peak, or increased sensitivity to lower or higher spatial frequencies (see group means in **Table S4** and detailed ANOVA results in **Figure S7**). Our observers trained on a 6-cpd Gabor, a challenging spatial frequency for amblyopes, but not close to their spatial frequency cutoff. A VPL study in which amblyopic observers showed increased sensitivity to higher spatial frequencies had observers train on a contrast sensitivity task near their individual SF cutoffs (Huang et al., 2008). Both groups also showed modest increases in the AULCSF, similar in magnitude to another VPL training study with amblyopic adults (Gu et al., 2020), convergent evidence of foveal contrast sensitivity improvements. Notably, the Attention group showed AULCSF gains in both eyes, whereas the Neutral group only showed gains in the amblyopic eye (although there was less room for improvement in the fellow eye) (**Figure 4**). In partial agreement with the qCSF, both groups showed similar improvement in amblyopic eye Pelli-Robson contrast sensitivity for letters, with less improvement for the fellow eye (**Figure 4** & **Table S3)**.

#### Improvements in logMar visual acuity

Amblyopic observers often experience concomitant improvements in visual acuity regardless of the VPL training task (reviews: Levi and Li, 2009; Astle et al, 2011; Hess et al., 2015; Levi, 2020; Rodan et al., 2020), but show the largest gains after training on contrast sensitivity-dependent tasks (review: Levi and Li, 2009). Given that visual acuity deficits are a classic diagnostic marker of amblyopia, we were particularly interested in measuring the interocular difference in observers’ visual acuity at the fovea. A VPL review of studies in amblyopic adults found an average amblyopic eye mean improvement of .17 logMar, with a third of participants achieving gains >.2 (Tsirlin et al., 2015). Notably, most of our observers, even those who may not have improved on the main task, improved in visual acuity: Almost 2 lines in the amblyopic eye in both groups, with smaller improvements for the fellow eye (less than half a line for the Neutral group and a little less than 1 line for the Attention group), which had less room for improvement (**Figure 4** & **Table S3**). The magnitude of these gains is similar to those found by other VPL studies that have directly trained amblyopic adults’ contrast-sensitivity (e.g., Polat et al., 2004; Zhou et al., 2006; Zhang et al., 2014). Remarkably, training reduced the interocular visual acuity difference, or amblyopic depth, by ∼50% across both groups. Whereas we do not have an explanation for this pattern of results, in agreement with (Liu et al., 2011), our findings suggest that the time-course or VPL is faster for acuity than contrast sensitivity in the amblyopic brain.

#### Improvements in stereoacuity

Given the controversy that monocular training while patching the fellow eye may promote interocular suppression over binocularity (Hess et al., 2015), it is encouraging that stereoacuity thresholds improved (decreased) in both groups (**Figure 4**, **Table S3** & **Figure S11** for individual scatter plots). This finding reflects changes to observers’ typical binocular vision even when not wearing an eye patch, consistent with evidence that latent binocularity remains in the amblyopic brain, and can be at least partially recovered given the right conditions (e.g., Gu et al., 2020; review: Hess et al., 2015). PL studies in amblyopia have found incidental improvements in stereoacuity (e.g., Zhang et al., 2014; reviews: Levi and Li, 2009; Hess et al., 2015; Levi, 2020; Rodan et al., 2020). Relevant to our study, ‘push-pull’ training protocols involve “exciting” the under responsive amblyopic eye with transient cues which actively inhibit the overbearing fellow eye, in attempts to restore the proper interocular weighting of visual processing for each eye (reviews: Hess et al., 2015; Levi et al., 2015). It is possible that our training protocol, and the use of transient precues, helped recover residual binocular functioning by simultaneously increasing the signal strength in the amblyopic eye and reducing the suppressive drive from the fellow eye to yield more optimal levels of binocular summation.

#### No meaningful changes in crowding

In contrast to the other tasks, the average critical spacing of crowding did not reliably change in either eye of both groups (**Figure 4**, **Table S3** & **Figure S12** for individual scatter plots). This helps rule out that improvements on other tasks in our study were simply due to procedural or motor learning, in which observers improve because they get more comfortable with the motor demands of the task, rather than attaining the longer-lasting changes in perceptual sensitivity or discriminability that define VPL (review: Sagi, 2011). We observed a significant correlation between observers’ improvements in crowding and stereoacuity, further support for the relation between these two visual tasks in amblyopic individuals (Song et al., 2014). We also found a correlation between improvements in crowding and visual acuity, although crowding and visual acuity are weakly linked in amblyopia (Pelli and Yiltiz, 2017); some observers may suffer from worse crowding but have near normal acuity, and vice versa (Song et al., 2014).

The mechanisms underlying crowding in amblyopic foveal vision may be more similar to crowding in the periphery of neurotypical adults, which has been proposed to result from an extended pooling stage following the feature detection stage (Hariharan et al., 2005). Critical spacing may be indicative of neural density (active neurons per square degree; Strappini et al., 2017). Thus, sparse neural density may underlie amblyopic crowding deficits, and may help explain why we did not see changes. Perceptual learning studies with amblyopes have shown that crowding can be reduced after practicing directly on a crowding task (e.g., Hussain et al., 2012), but training on our contrast sensitivity task did not transfer to crowding.

### Promoting transfer over specificity

A goal of visual training in amblyopia and other visual disorders is to maximize improvement in every way – across spatial locations, features, tasks, visual dimensions and eyes. Thus, our design included several aspects known to promote greater transfer of learning over specificity:

i. simply having exposed stimuli at the untrained diagonal in the main task during the practice, adaptive staircase and pre-test blocks may have facilitated transfer (e.g., Wang et al., 2012).
ii. training at two locations along one diagonal, rather than at one location, likely promoted generalization (e.g., Donovan et al., 2015; Donovan and Carrasco, 2018; Donovan et al., 2020).
iii. training on stimuli of differing difficulty (a higher and lower contrast) that were randomly intermixed within each block. Including some easier trials promotes location and orientation transfer (e.g., Ahissar and Hochstein, 1997; review: Seitz, 2017). Likewise, blocks consisting of several short staircases, which contain a higher proportion of easier trials, have shown to facilitate VPL transfer in neurotypical adults (e.g., Hung and Seitz, 2014).
iv. training on contrast-based tasks leads to greater degrees of improvement and transfer than training on acuity-based tasks in adults with amblyopia (reviews: Astle et al., 2011; Levi and Li, 2009).

There are many other known factors to influence VPL transfer, including sensory adaptation (Harris et al., 2012), task precision (Jeter et al., 2009), length of training (Jeter et al., 2010), and sensory uncertainty of stimulus features in visual search (Yashar and Denison, 2017). As all of those factors were kept constant across the two training groups, they cannot explain different patterns of VPL transfer.

### Individual variability in VPL

As often the case with perceptual learning studies (reviews: Hou et al., 2016; Dosher and Lu, 2017; Seitz, 2017; Maniglia and Seitz, 2018) and particularly with amblyopia (reviews: Levi and Li, 2009; Astle et al., 2011; Sengpiel, 2014; Tsirlin et al., 2015; Levi, 2020; Rodán et al., 2020), there was a substantial degree of individual variability in the magnitude of learning at the trained and untrained conditions. Among those who improved at the trained diagonal, there was variability in the degree of transfer to the untrained diagonal, whether none, partial or complete (i.e., equal learning at the trained and untrained diagonals), in both groups. Critically, the *percentage* of those observers that exhibited partial or complete transfer of VPL was significantly greater in the Attention group than the Neutral group. There was also significant variability in the degree of concomitant improvement to the complementary series of untrained tasks. The present findings are similar to other studies that have reported notable intra-individual variability in both the magnitude of overall VPL and degree of transfer, but find that reliable patterns emerge at the group level (e.g., Donovan et al., 2015, reviews: Levi and Li, 2009; Levi, 2020; Rodán et al., 2020).

#### Optimal amount of training

There is evidence that the number of training hours that it takes for an observer to achieve asymptotic performance directly scales with the severity of their deficit (Li et al. 2008). Thus, an observer with mild amblyopia may asymptote within ten training sessions, whereas a more severe observer may barely start performing above chance within the same number of sessions and practice blocks. Thus, a standard number of training sessions would not likely capture the full learning curves for each individual observer. This mismatch between an individual’s optimal number of training sessions and our standard of ten training sessions may partially explain why most observers exhibited significant VPL while a minority did not.

#### Training protocol

Our visual training task was relatively demanding; observers were required to discriminate a peripheral target while maintaining fixation within a relatively tight fixation window. If their eye moved outside of the window, the trial would be cancelled, and they would have to redo the trial at the end of the block. As some observers suffered from greater fixational instability, especially those with the strabismic (“lazy eye”) or mixed subtypes of amblyopia, not all could complete the same number of training blocks each day before becoming fatigued. In these few cases (5 out of 20), we had to spread the 80 training blocks across more than ten sessions. Importantly, the average number of training sessions per group did not significantly differ from each other.

Another source of variability was the different number of days that passed between training sessions. We strongly recommended that no more than two days pass between training sessions, as sleep consolidation and consistency have been shown to be critical factors in PL (Yotsumoto et al., 2009), but occasionally there were unavoidable scheduling conflicts. Fortunately, the average number of days that both groups of observers took to complete training did not significantly differ from each other (Neutral group: *M*=10.8±.49 sessions, Attention group: *M*=10.1±.10 sessions; *t*(18)=1.40, *p*=.18, spread across 21.3±1.10 days and 18.0±1.56 days, respectively; *t*(18)=1.30, *p*=.21). Other human factors out of the experimenters’ control, such as observers’ quality of sleep (common sources of variability in all VPL studies), may have contributed to the individual differences in VPL.

#### Modulatory role of amblyopic severity

Previous studies have found initial severity to be the biggest predictor of overall improvement (e.g., Yehezkel et al., 2016; reviews: Astle et al., 2011; Tsirlin et al., 2015; Rodán et al., 2020). Importantly, there were no pronounced differences in the composition of amblyopic severity (amblyopic eye visual acuity), and depth (interocular acuity difference) between the two groups (see **Table S1**). Across all visual tasks and observers, the amblyopic eye was worse than the fellow eye, providing confirmatory evidence for observers’ amblyopia diagnosis, and that the amblyopic contrast sensitivity deficit extends into the parafovea. As expected, all observers required more contrast in their amblyopic eye than fellow eye to complete the task with similar performance; but most importantly, the required stimulus contrast for each eye did not significantly differ between groups.

#### Amblyopia subtypes

It has been proposed that the various subtypes of amblyopia (i.e., anisometropic (refractive), strabismic (“lazy eye”), deprivational, or mixed) actually represent different visual populations, whose unique etiologies result in distinct constellations of visual deficits (McKee et al., 2003; Levi, 2006; Levi, 2013; Levi et al., 2015; Levi, 2020). For example, strabismic amblyopes are thought to have better contrast sensitivity but worse visual acuity than anisometropic amblyopes. However, a complementary statistical analysis (see ***Role of amblyopic subtype on main task performance***) revealed that the pattern of results for the main training task did not differ according to amblyopic subtype.

#### Differences in number of fixation breaks

Given their known issues with fixation stability, particularly for strabismic and mixed observers, one may wonder whether fixation breaks during the main task significantly affected observers’ performance (Verghese et al., 2019; Levi, 2020), as a higher preponderance would have taxed the system by extending the length of each training block. We did not observe a clear relation between the total number of times an observer broke fixation during a block and their overall performance (see **Role of fixation ability on main task performance** under **Results**). Further, observers’ total number of fixation breaks neither consistently changed across training, nor differed between the pre-test and post-test. The fact that we did not observe significant changes in all tasks, i.e., no reductions in the critical spacing of crowding, also supports the idea that task improvements were not attributed to alternative explanations such as improved gaze stability or accommodation. Together, the results indicate that individual differences in the ability to consistently fixate on the central cross cannot explain the observed variability in VPL across conditions and tasks.

### Selective attention in amblyopia

Some researchers have attributed differences in performance between amblyopic and neurotypical adults on a few higher-level psychophysical tasks –i.e., the attentional blink, numerosity estimation, and multiple object tracking–to an attention deficit in amblyopia. Critically, most have only inferred the role of attention in each of their tasks without directly manipulating it (see Roberts et al., 2016; Pham et al., 2018 for an expanded discussion). Recently, it has been hypothesized that increased fixation instability (particularly for strabismics) leads to higher rates of microsaccades, or fixational eye movements <1°, which results in an amblyopic deficit in selective attention (Verghese et al., 2019). However, there is controversy as to whether amblyopes reliably generate higher rates of microsaccades (see Discussion of Ramesh et al., 2020). Further, there is not a direct or causal correspondence between eye movements at any scale and voluntary or involuntary attention (review: Rolfs, 2009); neurotypical (review: Carrasco, 2011) and amblyopic (e.g., Roberts et al., 2016; Ramesh et al., 2020), and neurotypical observers can selectively attend within the high-acuity foveola without fixational eye movements and even with retinal stabilization (Poletti et al., 2017).

There is evidence that the neural mechanisms and correlates of voluntary attention differ in the amblyopic brain (Hou et al., 2016; Pham et al., 2018; Mortazavi et al., 2020). A study measuring the full contrast-response functions of amblyopic monkeys for a motion direction discrimination task found that, unlike the fellow eyes which exhibited only contrast gain, the amblyopic eyes showed both contrast and response gain, perhaps in a compensatory fashion (Pham et al., 2018). Steady-state visual evoked potentials while strabismic adults were cued to voluntarily attend one of two hemifields to detect a contrast increment in a peripheral grating revealed reduced attentional modulation of neural activity in striate and extrastriate cortex corresponding to the amblyopic eye, but only in the extrastriate cortex corresponding to the fellow eye (Hou et al., 2016). In a more recent study, amblyopic adults showed higher discriminability for the cued than uncued hemifield, but event-related potentials corresponding to the amblyopic eye had delayed latencies and reduced peak amplitudes in components associated with various stages of the visual processing hierarchy (Mortazavi et al., 2020).

The few studies employing precues to directly manipulate covert attention in amblyopia show that amblyopic adults (Sharma et al., 2000; Hou et al., 2016; Roberts et al., 2016; Mortazavi et al., 2020), children (Ramesh et al., 2020) and monkeys (Pham et al., 2018) benefit when spatially cued to voluntarily (Sharma et al., 2000; Roberts et al., 2016; Pham et al., 2018; Ramesh et al., 2020) and involuntarily (Roberts et al., 2016) attend to a task relevant spatial location and show a behavioral cost when their attention is shifted elsewhere (Sharma et al., 2000; Roberts et al., 2016). Covert selective spatial attention is behaviorally functionally intact in amblyopia, even if through alternative and potentially compensatory neural mechanisms. Given that we found differences in VPL transfer at post-test between our two groups, whose training only differed in training with or without attention, the current study provides further evidence that the adult amblyopic brain can effectively use a spatial cue during perceptual training to induce meaningful changes in perceptual processing.

In the present study we employed peripheral exogenous precues during an orientation discrimination task. All but one of the studies isolating the role of attention in amblyopia have explicitly manipulated *voluntary* attention. The only study with exogenous attention found that cues adjacent to the target location benefited orientation discrimination whereas those far away disadvantaged performance. These effects were as pronounced when observers used their amblyopic eye as when they used their fellow eye, and also as pronounced as the corresponding effects of neurotypical age- and gender-matched controls (Roberts et al., 2016). The effectiveness of transient cues presented exclusively to the weaker eye during push-pull binocular training programs for re-calibrating interocular balance (Levi et al. 2015) also suggests that the amblyopic eye robustly responds to bottom-up or stimulus-driven (exogenous attention) cues. More research should be done to better characterize the behavioral effects and largely unknown neural correlates of exogenous and endogenous attention in amblyopia using different visual tasks (e.g., visual search, the attentional blink paradigm, motion object tracking), and under conditions of increasing task difficulty and/or greater attentional load.

### How attention may generalize learning to untrained locations

From a mechanistic standpoint, modeling studies have revealed substantial signal attenuation, increased additive internal noise, and weaker perceptual templates in the amblyopic brain (reviews: Levi, 2013; Hess et al., 2015; Levi, 2020). Thus, as in other amblyopia VPL studies (e.g., Li and Levi, 2008), visual training may have acted to reduce this elevated internal noise and retune observer’s perceptual templates.

There are several conceptual (e.g., Maniglia and Seitz, 2018) and computational models of VPL (review: Dosher and Lu, 2017). Researchers have speculated that attention plays a critical role, but most have discussed it in terms of task relevance or in selectively “gating” what gets learned (Dosher and Lu, 2017; Maniglia and Seitz, 2018), and none has explicitly modeled the role of spatial attention on VPL. Although no current theory or model of VPL can explain why or how exogenous attention generalizes VPL to untrained spatial locations in the amblyopic brain, below we highlight how some models may relate to our findings:

*A rule-based learning model* suggests that VPL primarily involves learning rules for performing the task efficiently, and that specificity is a consequence of an inability to link signals from early visual cortex that represent untrained stimuli to the learned rule scheme (e.g., Zhang et al., 2014). This model cannot account for our findings, as the model predicts that: (a) for transfer to occur, the rule scheme must be learned before exposure to untrained locations or features. But for both groups, we exposed the untrained diagonal only during the pre-test, not during training; (b) transfer is prevented at unstimulated locations because they are suppressed. Accordingly, the Attention group should show more specificity, not transfer as we observed, because exogenous spatial attention decreases neural activity at unattended locations (e.g., Herrmann et al., 2010; Wang et al., 2015; Dugue et al., 2020; Fernández and Carrasco, 2020).

The *Reverse hierarchy theory* postulates that VPL is a top-down guided process that begins with higher-level regions and progresses to lower-level sensory regions with more stimulus repetitions and/or challenging perceptual tasks; location-specific learning is associated with plasticity in early visual regions and is more likely to occur for difficult tasks with noisy sensory signals (Ahissar et al., 2009). According to this framework, we speculate that location transfer in the Attention group could have resulted from exogenous attention improving the target’s signal-to-noise ratio and increasing its visibility, thereby making the task easier during training. Attention-induced modification would thus be more likely to occur in higher-level decision-making areas, e.g., lateral intraparietal cortex (LIP), which contain broadly tuned, bilateral receptive fields, enabling learning effects to transfer to untrained locations as far as in the opposite hemifield (Sagi et al., 2011; Dosher and Lu, 2017).

The *Integrated Reweighting Theory* explains transfer across retinotopic locations by incorporating higher-level, location-independent representations with lower-level, location-specific representations that are both dynamically modified in VPL (review: Dosher and Lu, 2017). According to this theory, specificity of VPL to the trained diagonal in the Neutral group would likely be due to the reweighting, or enhanced readout, of the lower-level, location-specific representations. Transfer in the Attention group would likely be due to reweighting of higher-level, location-invariant representations of the decision stage. Currently, the role of selective attention is not explicitly modeled in any VPL reweighting model. Future computational modeling studies may illuminate the specific mechanism(s) through which attention may promote location transfer.

### Where are the neural correlates of VPL in amblyopia?

VPL that is specific to locations and features is often considered to take place in primary visual cortex (reviews: Sagi, 2011; Seitz, 2017), as V1 neurons respond to precise retinal locations and simple visual features. The finding that our observers demonstrated complete interocular transfer of VPL suggests that the neuronal correlates of this learning most likely reside in visual cortex, where binocular convergence first occurs. But given that there are fewer binocularly driven neurons (reviews: Kiorpes and Daw, 2018; Levi, 2020) and abnormal patterns of binocular interactions in amblyopic V1 (reviews: Joly and Franko, 2014; Hess et al., 2015; Levi et al., 2015), VPL could have occurred either at primary visual cortex or downstream in higher-order vision or decision-making areas. Our findings suggest that attending to the target location during training may have increased neural activity in occipital cortex (e.g., Liu et al., 2005; Herrmann et al., 2010; Wang et al., 2015; Dugué et al., 2020; Fernández and Carrasco, 2020) and altered the distribution of VPL changes to be biased towards higher brain areas whose responses are less tuned to specific spatial locations, features and/or eyes. VPL for any task is widespread, likely involving several areas, and the pattern of VPL changes and their neural correlates may differ according to the training protocol and task type and difficulty (e.g., Jia et al., 2020; reviews: Gilbert and Li, 2012; Dosher and Lu, 2017; Seitz, 2017; Maniglia and Seitz, 2018).

### Translational implications for rehabilitation of visual disorders

With early detection and compliance, there are several treatments for children with amblyopia (e.g., refractive correction, surgical realignment of the eyes, patching) that largely mitigate the most severe symptoms from persisting into adulthood (reviews: Astle et al., 2011; Sengpiel, 2014; Levi, 2020). In contrast, treatment options for adults have been more limited. However, VPL studies have advanced our understanding of the impressive remaining neuroplasticity in the (a)typically developed adult brain and identified visual training as a promising rehabilitative tool. Perceptual training improves visual performance in individuals with amblyopia (reviews: Levi and Li, 2009; Astle et al., 2011; Sengpiel, 2014; Levi et al., 2015; Tsirlin et al., 2015; Levi, 2020; Rodán et al., 2020), visual acuity in presbyopia (Polat et al., 2012), visual acuity and contrast sensitivity in individuals with optical aberrations (Schoups and Orban, 1996), and visual motion discrimination in people with V1 damage (Cavanaugh et al., 2015; Cavanaugh et al., 2019). But the prognoses for most visual disorders remain poor and achieving any improvements in daily vision takes a long time.

Most current VPL protocols that promote location transfer in neurotypical adults (e.g., Szpiro et al., 2014; Wang et al., 2014; Hung and Seitz, 2018) are limited in their practicality for clinical populations; they would demand a lot of effort from an already bioenergetically taxed system, and patients would have to practice more than one task to achieve maximal transfer. Here we present a simple and elegant possibility for improving visual training efficiency –training with exogenous attention (Donovan et al., 2015; Szpiro and Carrasco, 2015; Donovan et al., 2020; Nguyen et al., 2020). Exogenous attention cues could easily be incorporated into many types of training treatments currently being offered to individuals with amblyopia, from relatively simple perceptual learning tasks while patching to more complex dichoptically-presented videogames. These attention cues would enhance VPL at little-to-no monetary cost or additional effort from the patients by facilitating its generalization to untrained spatial locations, which would be a major benefit. This study complements studies that have shown the practical benefits of videogame play for improving amblyopic and neurotypical vision on a broad range of dimensions (Levi et al., 2015; Tsirlin et al., 2015; Bavelier & Green, 2019; Levi, 2020), as it provides insight into the specific role involuntary spatial attention likely plays in facilitating more generalized learning, including across visual space.

### Conclusions

This study reveals that training on a discrimination task in the parafovea leads to improvements in foveal contrast sensitivity, acuity and stereoacuity in adults with amblyopia. Notably, exogenous attention generalizes the effects of perceptual training to untrained spatial locations.

### Limitations of the study

Due to the COVID lockdown, we were unable to collect the post-test data for one observer in the Attention group for all tasks in the visual battery, nor add more observers per group. We were also unable to bring observers back for a 6-month follow-up to see how much of the learning was retained after an extended period, as we had originally planned. Given our relatively modest effect sizes and high individual variability, future replications of our study with more observers will be important to verify the robustness of our findings.

## Acknowledgments

This research was supported by National Eye Institute RO1-EY027401 to M.C., National Eye Institute T32-EY007136 to New York University, National Science Foundation Division of Graduate Education DGE-1342536 to M.R., and the New York University Graduate School of Arts & Sciences Dean’s Dissertation Fellowship to M.R. We thank Julia Payne and Sanjana Manjunath for their help with data collection, Antonio Fernández, Shao-Chin Hung, Michael Jigo, Caroline Myers, Angela Shen and other members of the Carrasco Lab, as well as Sarit Szpiro, Amit Yashar, Roozbeh Kiani, Lynne Kiorpes, Zhong-Ling Lu and Denis Pelli for helpful comments.

## Author Contributions

M.C. and M.R. designed the experiments, interpreted the data and wrote the manuscript. M.R. programmed the experiment and collected and analyzed the data.

## STAR METHODS

### RESOURCE AVAILABILITY

#### Lead contact

Further information and requests should be directed to the lead contact, Mariel Roberts.

#### Materials availability

This study did not generate new unique reagents.

#### Data and code availability

- All data files for the trained and untrained tasks are freely available on Mendeley at (http://dx.doi.org/[10.17632/g2fnm9fv3h.1]).
- All original experimental code is freely available via an open-access repository (https://github.com/CarrascoLab/Exo-Att-PL-Amblyopia).
- Any additional information required to reanalyze the data reported in this paper is available from the lead contact upon request.

### EXPERIMENTAL MODEL AND SUBJECT DETAILS

The observers who had been prescribed refractive correction from their personal eye doctor were required to wear either the exact same pair of glasses or contacts that they would typically wear throughout the entire study. Before training, we confirmed that observers exhibited a ≥ two-line interocular difference in EDTRS logMar acuity (also known as amblyopic depth), in which each letter on the eye chart equals .02 and a full line equals 0.1, 0 is equivalent to 20/20 Snellen acuity, and a higher score is worse (amblyopic depth: Neutral group *M*=.546±.062, Attention group *M*=.594±.088; *t*(18)=-.462, *p*=.650). Five observers who qualified decided not to participate and complete the pre-training phase (days 1-4), and one observer per group completed less than half of the training days. Ultimately, twenty adults with amblyopia and no other visual disorder or disease were included in the study (see **Table S1** for demographic and clinical information). All experimental procedures were in agreement with the Helsinki declaration and approved by the New York University Institutional Review Board. All observers were naïve to the experimental hypotheses and signed a written consent.

### METHOD DETAILS

#### Apparatus & setup

All computer-based tasks shared the same setup. Experiments were run from a 21.5” Apple iMac 3.06 GHz Intel Core 2 Duo Desktop using MATLAB R2014b (MathWorks, USA) in conjunction with the MGL toolbox (http://justingardner.net/mgl). They were presented at a viewing distance of 114 cm on a 40x30 cm IBM-P260 CRT monitor (1280x960 pix resolution, 90- Hz refresh rate), calibrated and linearized using a Photo Research PR-650 SpectraScan Colorimeter.

Eye movements were monocularly monitored using an EyeLink 1000 Desktop Mount eye tracker (SR Research, Canada) with a 1000-Hz sampling rate. We used a 5-point display to calibrate the eyetracker at the beginning of every block and within blocks as needed to minimize fixation breaks. Observers’ logMar visual acuity was measured using the 2M Original Series EDTRS Chart R (Precision Optics Ltd., CO)(Bailey and Lovie-Kitchen, 2013).

#### Procedure

There were three phases to the study, which took place across at least 17 different sessions (**Figure 1b**). During the pre-training phase (Phase I; days 1-4), we prescreened observers for qualification in the study (see **Observers**) and measured their abilities on a variety of visual tasks. During Phase II, observers performed the same training task for ∼10 sessions while being presented with either a neutral (Neutral group) or valid (Attention group) attention precue. The three final sessions of the post-training phase (Phase III; days 15-17) were identical to those of Phase 1, except these sessions did not include the adaptive staircase session for measuring contrast thresholds from day 2 and were administered in reverse order.

##### Phases I & III: Pre-training & Post-training

Day 1 consisted of a prescreening session in which we determined whether observers fulfilled our inclusion criteria (see **Observers**) and obtained a baseline measure of their abilities on a curated selection of visual tests known to be impaired in amblyopia: foveal letter contrast sensitivity using a Pelli-Robson chart, logMar visual acuity, crowding, and stereoacuity (see **Supplemental Information** for details). After obtaining consent and having participants fill out a demographic survey, we conducted a full oral interview of their visual history. The protocols for days 1 and 17 were identical from this point on. We measured observers’ foveal logMar visual acuity and Pelli-Robson letter contrast sensitivity by having them read from eye charts (see **Apparatus & setup**) monocularly (fellow eye always first) with the other eye patched, from 2m away in a consistently brightly lit room. All remaining tasks were administered in a darkened and sound-attenuated room. Observers used a forehead and chin rest to ensure head stabilization and a constant viewing distance. On average, the prescreening session on day 1 took 2 hours and the post-training session on day 17 took 1.5 hours.

For all computer-based tasks administered during days 2 through 16, if observers moved their eyes ≥1° from the fixation cross, the text ‘Please fixate’ would appear onscreen and that trial would be replaced at the end of the block. On days 2 and 16, we estimated observers’ foveal broadband contrast sensitivity in their fellow and amblyopic eyes separately using the quick Contrast Sensitivity Function (qCSF) procedure (see **Supplemental Information** for details).

On day 3, to account for procedural learning, observers first performed a simple two-alternative forced-choice (2AFC) orientation discrimination task (2-cpd, 64% contrast, 3.2°-wide Gabor tilted ±5° from horizontal) at the fovea for ≥40 trials. Next, they completed 1-2 practice blocks of 80 trials along each diagonal. Two 3.2°-wide 6-cpd Gabors (64% contrast) simultaneously appeared at 4° eccentricity along one of the diagonals. The observer had to report the orientation (±4° from horizontal) of the target indicated by a central response cue.

Observers then completed an adaptive staircase procedure to estimate the monocular contrast thresholds required for the amblyopic and fellow eye to perform the main training task with similar accuracy at pre-test along the same diagonal to which they were randomly assigned to train (upper-right and lower-left quadrants or upper-left and lower-right quadrants). Observers first completed four 40-trial blocks with their fellow eye (with amblyopic eye patched), and after a break, they completed four blocks with their amblyopic eye (with fellow eye patched). Each block contained two 20-trial interleaved staircases to estimate the contrast thresholds corresponding to 75% and 88% accuracy. On each trial of a staircase, the contrast of the Gabors adjusted according to best PEST —an adaptive staircase procedure as implemented in the Palamedes toolbox (Prins and Kingdom, 2009). A uniform prior was used for the first block of each staircase, and starting from the second block, the posterior of the previous block was used as the prior for the next. The threshold estimates from the last three blocks were averaged to estimate the individual observers’ contrast thresholds for a given eye. Observers were only presented with neutral attention cues (see **Main task stimuli**) during the adaptive staircase procedure. On average this session took 1.5-2 hours.

Days 4 and 15 were the pre-test and post-test sessions for the main training task. Before running the main task pre-test in each eye, we conducted one more block of the adaptive staircase procedure to assess whether the two contrast threshold estimates were similar to those in the prior session. If there was a large discrepancy (perhaps due to some short-term consolidation of learning), the experimenter would manually adjust the stimulus contrast levels for each eye according to their best judgment to better capture the dynamic range of the observer and make the task challenging enough that they would have plenty of room for improvement with training (i.e., performed with a d’ around 1 overall in each eye). These stimulus contrast levels were then held constant throughout the remainder of the study, including the post-test. Throughout each block of the main task, the contrast of the stimuli randomly varied between these two levels so that observers performed a mix of harder and easier trials. Given that we did not find significant differences in the results between contrast levels, all data and analyses reported correspond to performance collapsed across them.

During the pre-test and post-test sessions, we measured baseline performance on the main task in four conditions while observers were only exposed to neutral cues, in this order: (1) fellow eye along the upper-right and lower-left diagonal, (2) amblyopic eye along the upper-right and lower-left diagonal, (3) fellow eye along the upper-left and lower-right diagonal, and (4) amblyopic eye along the upper-left and lower-right diagonal. On average these pre-test and post-test sessions took 1.5-2 hours.

##### Phase II: Training

Observers completed 6,400 trials total: approximately eight 80-trial blocks during each of ∼10 training sessions. We hypothesized that this training protocol would be sufficient to result in significant VPL for the majority of observers with amblyopia, but not so much that it would promote specificity, as has been shown in neurotypical adults (Jeter et al., 2010).

During each 1-hour training session, observers completed as many 80-trial blocks of the main 2AFC orientation task (see **Main task trial sequence**) as possible using their amblyopic eye with their fellow eye patched. During training sessions, observers took 30-second breaks halfway through each block (a neutral auditory tone indicated when they should open their eyes), paused between blocks, and had a 5-minute break halfway through to minimize oculomotor fatigue and retinal adaptation. Each observer was randomly assigned to train along one of the two diagonals (five observers trained on each diagonal within each group) while being exposed to only neutral cues (Neutral group; *n*=10), which spread an observer’s attention evenly across both potential target locations, or 100% valid peripheral cues (Attention group; *n*=10), which rapidly and automatically (i.e., requiring no additional effort or even awareness of the cue from the observer) focuses their attention to the upcoming target location. We used only valid peripheral cues, and not invalid cues, which would direct attention to the nontarget location and decrease performance (e.g., Pestilli & Carrasco, 2005; Barbot et al., 2012, Fernández & Carrasco, 2020; Dugué et al 2020). Cue validity does not affect the magnitude of either the benefit or cost of exogenous attention on performance during classic spatial cueing experiments (Giordano et al., 2009; review: Carrasco, 2011).

#### Main task stimuli

Observers were asked to fixate on a black cross (1° across) at the center of the screen throughout the trial (**Figure 1a**). The target and distractor stimuli were both 3.2°-wide, 6-cpd Gabor patches (contrast-defined sinusoidal gratings embedded in a Gaussian envelope, σ=0.46°), randomly and independently tilted ±4° from horizontal, centered at 4° eccentricity along the diagonals, and with the same mean luminance as the uniform grey background (26 cd/m^2^). The neutral precue consisted of four 0.16° white dots surrounding the fixation cross and centered 0.5° away from its edges. The peripheral exogenous attention precue was identical to the neutral cue but centered around and placed 0.5° from the edge of the target in the periphery, to prevent masking. The response cue (a 0.8°-long white line starting 0.8° from the center of the screen) indicated the target location by pointing to one stimulus location (matching the peripheral attention precue with 100% validity). Similar central neutral and peripheral valid cues have been used in several studies of exogenous attention (e.g., Barbot et al., 2012; Szpiro & Carrasco, 2015; Roberts et al., 2016; Donovan et al., 2015, 2020; Li et al., 2021). Furthermore, the magnitude of the effect is the same with the central neutral cue employed here and with a distributed cue (cues adjacent to all potential target locations; e.g., Talgar et al., 2004; Carrasco et al., 2002).

#### Main task trial sequence

Observers monocularly (with the other eye patched) performed a 2AFC orientation discrimination task while their exogenous spatial attention was manipulated via presentation of a brief valid peripheral precue (**Figure 1a**). After centrally fixating (256 ms), the precue was presented (67 ms), after which there was a brief interstimulus interval (ISI; 67 ms) during which the screen was blank. The target and distractor Gabor stimuli then appeared at isoeccentric locations along one of the diagonals concurrently with the response cue for 123 ms. This 134-ms stimulus-onset-asynchrony between the onset of the precue and onset of the stimuli enabled us to assess the attentional effects of the exogenous cue while preventing any voluntary deployment of attention (review: Carrasco, 2011). After the Gabors disappeared, the response cue stayed throughout the remainder of the trial. 78-ms after the stimuli were removed, the fixation cross turned white and an auditory tone indicated the beginning of the response window, in which observers had to report the target orientation (counterclockwise or clockwise relative to horizontal) using one of two keyboard presses (‘1’ for counterclockwise, ‘2’ for clockwise) with their right hand. On every trial, observers were encouraged to respond as accurately as possible within a 5-s response window. Observer response terminated the trial, after which there was a mandatory 1000-ms intertrial interval. Auditory feedback was provided at the end of each trial and visual feedback indicating observers’ accuracy and number of fixation breaks was presented at the end of each block.

#### Untrained task procedures

##### qCSF

We incorporated the qCSF into a 4AFC orientation discrimination task (**Figure S6**, left side). The 5.6°-wide target stimulus could be any combination of parameters from a 2D array of spatial frequencies (12 levels ranging from 0.25- to 20-cpd) and contrasts (60 levels ranging from 1% to 100%; **Figure S6**, right side). After at least one practice block in which they binocularly performed the task above chance using an easier (higher contrast range) stimulus set, observers completed three 70-trial blocks with each eye while the other was patched. Observers always started with their fellow eye and alternated eyes every block. If observers moved their eyes more than 1° from the fixation cross, the text ‘Please fixate’ would appear onscreen and that trial would be cancelled and added to the end of the block. On average the qCSF session took 75 min.

##### Pelli-Robson contrast sensitivity

Log contrast sensitivity thresholds for letters were measured using one of two versions of the Pelli-Robson Contrast Sensitivity Chart (Pelli et al., 1988). Participants monocularly (with the other eye patched) read one version with their fellow eye and then the other version with their amblyopic eye. Pelli-Robson scores were graded as the log contrast sensitivity threshold corresponding to the last set of letter triplets in which they identified two letters correctly, reading from top left to bottom right (Pelli et al., 1988), with higher scores indicating greater contrast sensitivity.

##### Stereoacuity thresholds

Stereoacuity thresholds were measured using the ASTEROID software program (Vancleef et al., 2019). The task was presented on an Android 3D stereotablet presented at eye-level and parallel to the observer’s head 25 cm away, held stable by a tablet stand whose base was clamped to the table. Observers performed the task in a darkened and sound-attenuated room with their head in a forehead and chin rest to ensure head stabilization. Observers were instructed to keep both eyes open and detect an illusory 3D square among four possible locations, then tap the corresponding quadrant of the screen. Observers first performed a short practice block of 20 trials, then completed two different adaptive Bayesian staircase blocks of 60 trials each. We averaged these two blocks to estimate an observer’s stereoacuity threshold.

##### Critical spacing of crowding

We measured observers’ critical spacing (degrees of visual angle required to reliably and accurately discriminate a central number from two flanking distractor numbers to the left and right) at the fovea using the Pelli Critical Spacing toolbox (Pelli et al., 2016) for MATLAB. The task was presented on a 13-inch 2017 MacBook Air using MacOS Mojave in a dark and sound-attenuated room, with observers’ heads stabilized by a forehead and chin rest. The stimulus display screen was flipped, then presented onto a mirror 114 cm away from the observer to simulate presenting the task at 228 cm, the necessary distance for displaying letters of a sufficient range of sizes to measure small thresholds. The experimenter input the observer’s verbal responses manually using an external keyboard attached to the laptop. After binocularly performing above chance in a 10-trial practice block, observers completed a 20-trial adaptive staircase three times in a row with each eye (with the other eye patched). We averaged the estimates from all three staircases to estimate the critical spacing of crowding for each eye.

### QUANTIFICATION AND STATISTICAL ANALYSIS

We measured changes in d’ as the primary dependent variable to index changes in observers’ perceptual sensitivity independent of their response criterion. d’ was calculated as the z-score of the proportion of hits (arbitrarily chosen as when target was counterclockwise (CCW), observer reported CCW) minus the z-score of the proportion of false alarms (target was CCW, observer reported clockwise (CW); e.g., Herrmann et al., 2010; Dugué et al., 2016; Pham et al., 2018; Dugué et al., 2020; Fernández and Carrasco, 2020; Jigo and Carrasco, 2020). To avoid infinite values, we added 0.5 to the number of hits, misses, correct rejections and false alarms and 1 to the number of “signal present” (i.e., CCW target) and “signal absent” (i.e., CW target) trials prior to computing d’ (Brown and White, 2005). To account for between-subject differences in pre-test performance (in spite of doing our best to equate initial performance by adjusting stimulus contrast) and to more fairly compare learning across different tasks with different units of measurement, we conducted complementary analyses for most tasks with our own performance measure – normalized change– which we defined as the difference between post-test and pre-test performance divided by the sum of their values (except for logMar acuity, for which we simply calculated pre-test minus post-test as they are log values). A positive normalized change indicated *improvement*. Thus, for those tasks in which a lower score is better (e.g., critical spacing of crowding), the change was calculated as pre-test minus post-test performance.

We designed our task to evaluate changes in orientation sensitivity (d’). Observers were encouraged to be as accurate as possible. Median RTs for correct trials served as a secondary variable to assess potential speed-accuracy tradeoffs. RTs were measured relative to the simultaneous onset of the Gabor stimuli and response cue (**Figure 1a**), which remained onscreen for 123 ms. The Gabors disappeared, leaving only the response cue, and observers had to wait an additional 78 ms before they were allowed to respond within a 5-s window, to further encourage observers to be as accurate rather than as fast as possible. In our experimental design, as in many tasks that measure RT as the primary dependent variable (e.g., Wang et al., 2015), it was impossible to distinguish the degree to which reaction time (RT) differences reflected changes in speed of processing, discriminability, response criterion, and/or motor learning (e.g., Carrasco and McElree, 2001; Roberts et al., 2017).

We conducted all ANOVAs in R with ezANOVA using the Type-III sums of squares approach. We always included Observer as a random factor, Eye (amblyopic, fellow), Diagonal (trained, untrained) and Session (pre-test, post-test) as fixed within-subjects factors, and Group (Neutral, Attention) as a fixed between-subjects factor. Planned comparisons were two-tailed paired t-tests, and the Benjamini-Hochberg adjustment (Benjamini and Hochberg, 1995) was applied to correct for multiple comparisons as it controls for the false discovery rate. All reports of variability represent ±1 standard error of the mean.

## SUPPLEMENTAL INFORMATION

**Table S1.** related to **STAR Methods.** Demographic and clinical information for all observers. Precluded observers from the main analysis (see **Results**) are shaded in grey.

| Participant | Gender | Age | Amblyopic eye (L/R) | Depth<sup>1</sup> | Severity<sup>2</sup> | Subtype<sup>3</sup> | logMAR acuity<sup>4</sup> | Refractive correction<sup>5</sup> | Past treatment |
| --- | --- | --- | --- | --- | --- | --- | --- | --- | --- |
| **Neutral group** |  |  |  |  |  |  |  |  |  |
| 1 | F | 29 | L | 0.78 | S | S | 0.84 | none | patching, glasses, visual training for 2 yrs |
| 2 | F | 22 | R | 0.60 | S | M | 0.62 | glasses; OD: +3.00, OS: +2.00 | patching |
| 3 | F | 21 | R | 0.80 | S | A | 0.72 | contacts; OD: +6.5 -1.00 × 160, OS: +2.00 -0.25 × 100 | patching, eyedrops, corrective lenses |
| 4 | F | 27 | L | 0.60 | MO | M | 0.52 | glasses; OD: -5.00 -0.75 × 160, OS: -7.5 -3.75 × 007 | patching, corrective surgery, corrective lenses |
| 5 | M | 24 | R | 0.74 | MO | M | 0.54 | glasses; OD: +2.75 -0.5 × 105, OS: -0.5 -0.25 × 175 | patching, corrective lenses |
| 6 | M | 24 | R | 0.30 | MO | M | 0.32 | glasses; OD: +4.25 -3.25 × 179, OS: +2.75 -2.25 × 009 | noncompliant patching, corrective lenses |
| 7 | F | 22 | L | 0.48 | MO | A | 0.46 | contacts; OD: -4.00 -1.75 × 160, OS: -1.50 -0.75 × 180 | glasses |
| 8 | M | 21 | L | 0.52 | MO | M | 0.56 | contacts; OD: -7.25 BC 8.4 DIA 13.8, OS: -6.75 BC 8.4 DIA 13.8 | noncompliant patching, glasses |
| 9 | F | 21 | R | 0.24 | MI | S | 0.18 | contacts; OD: +3.75, OS: +3.75 | patching, stereo training, corrective lenses |
| 10 | F | 19 | L | 0.40 | MO | S | 0.52 | glasses; OD: 3.00, OS: -3.75 | patching, eye strengthening exercises, corrective lenses |
| **Totals** | **M: 3**<br>**F: 7** | **M = 23.0±.966** | **L: 5**<br>**R: 5** | **M = .546±.062** | **S: 3**<br>**MO: 6**<br>**MI: 1** | **S: 3**<br>**A: 2**<br>**M: 5** | **M = .528±.059** | **none: 1**<br>**glasses: 5**<br>**contacts: 4** | **patching: 7**<br>**corrective surgery: 1**<br>**corrective lenses: 10**<br>**visual training exercises: 3** |
| **Attention group** |  |  |  |  |  |  |  |  |  |
| 1 | F | 21 | L | 0.42 | MO | A | 0.54 | contacts; prescription unavailable | none |
| 2 | F | 20 | L | 0.60 | MO | S | 0.52 | contacts; prescription unavailable | patching |
| 3 | F | 22 | L | 1.02 | S | M | 0.94 | glasses; prescription unavailable | patching |
| 4 | F | 19 | R | 0.66 | MO | A | 0.54 | contacts; OD: 8.5, OS: -2.5 | none |
| 5 | M | 22 | L | 0.56 | MO | S | 0.52 | none | patching, noncompliant eye drops, glasses |
| 6 | M | 20 | R | 0.44 | MO | S | 0.36 | none | stereo training |
| 7 | M | 33 | R | 0.22 | MI | M | 0.12 | glasses; OD: +3.5 -2.5 × 180, OS: -4.00 -1.5 × 180 | patching, glasses |
| 8 | M | 29 | L | 0.80 | S | A | 0.82 | none | noncompliant patch and eye strengthening exercises |
| 9 | F | 19 | L | 0.26 | MI | M | 0.22 | contacts; OD: -0.50 -0.25 × 62, OS: +2.00 | noncompliant patching, refractive correction |
| 10 | M | 37 | R | 0.46 | MO | S | 0.42 | glasses; OD: +3.75 -1.25 × 175, OS: +3.00 -1.50 × 018 | patching, glasses |
| **Totals** | **M: 5**<br>**F: 5** | **M = 24.2 ± 2.037** | **L: 6**<br>**R: 4** | **M = .594±.088** | **S: 2**<br>**MO: 6**<br>**MI: 2** | **S: 4**<br>**A: 3**<br>**M: 3** | **M = .509±.087** | **none: 3**<br>**glasses: 3**<br>**contacts: 4** | **patching: 6**<br>**corrective surgery: 0**<br>**corrective lenses: 7**<br>**visual training exercises: 1** |

<sup>1</sup>Depth = (LogMAR visual acuity (VA) amblyopic eye - LogMAR VA fellow eye) at pre-test while wearing prescribed refractive correction (if any); as calculated in Popple & Levi, 2008.
<sup>2</sup>Amblyopic severity at pre-test (according to amblyopic eye logMAR acuity; Stewart et al., 2005): MI = mild (0.1-0.3 logMar), MO = moderate (>0.3-0.6 logMar), S = severe (>0.6 logMar)
<sup>3</sup>S = strabismic, A = anisometropic, M = mixed
<sup>4</sup>Amblyopic eye logMAR acuity at pre-test
<sup>5</sup>Type of refractive correction and prescription (if available) that observer wore throughout study; none = observer did not wear refractive correction in or out of the study as they did not find it useful

**Figure S1.**
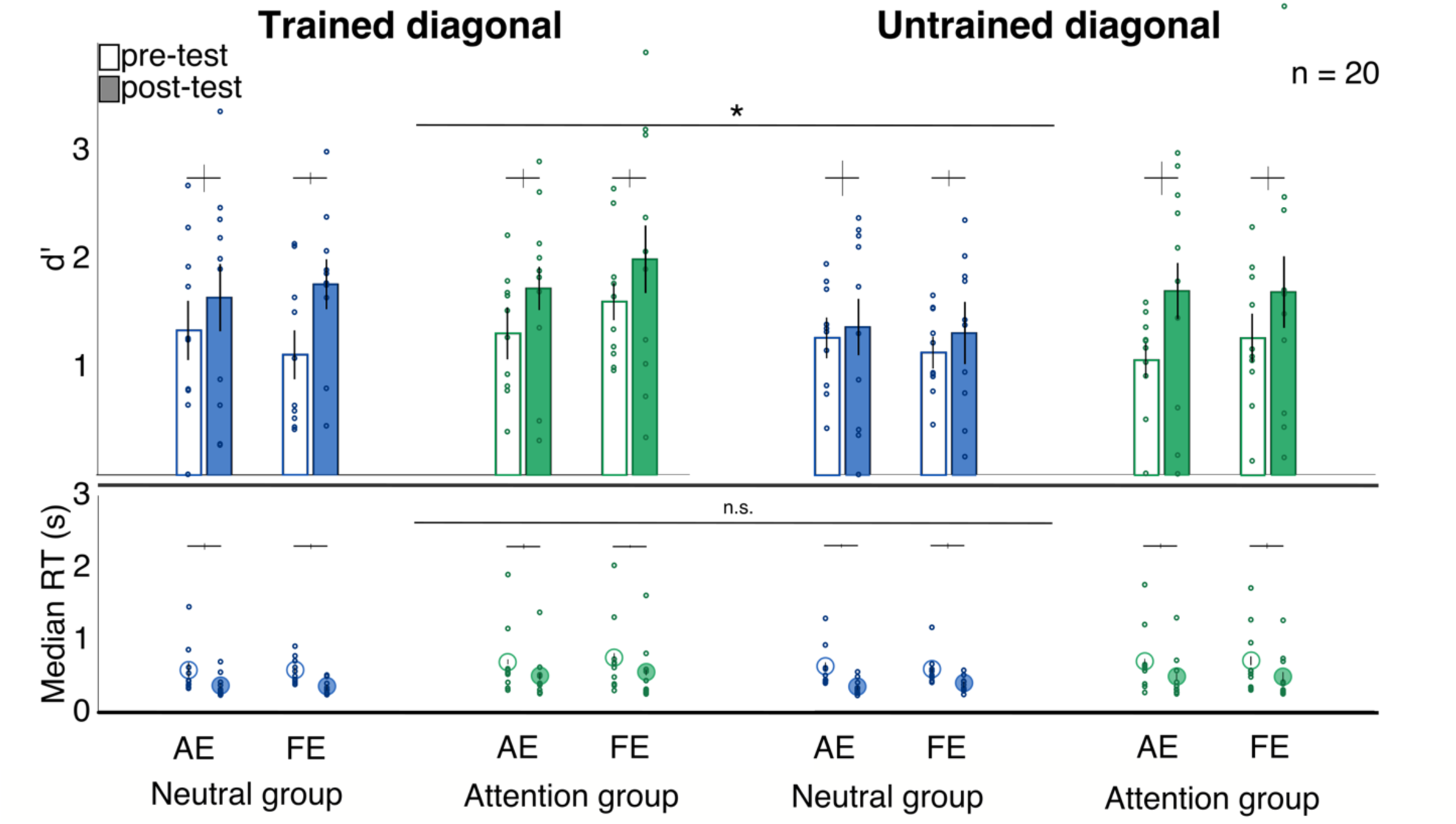
related to **Figure S2** & **S3.** Performance for all observers on the main training task at pre-test and post-test, split by diagonal, group and eye. Dots indicate the performance of individual observers. AE = amblyopic eye; FE = fellow eye. Error bars represent ±1 within-subject standard error (Morey, 2008). \**p*<.05.

**Figure S2.**
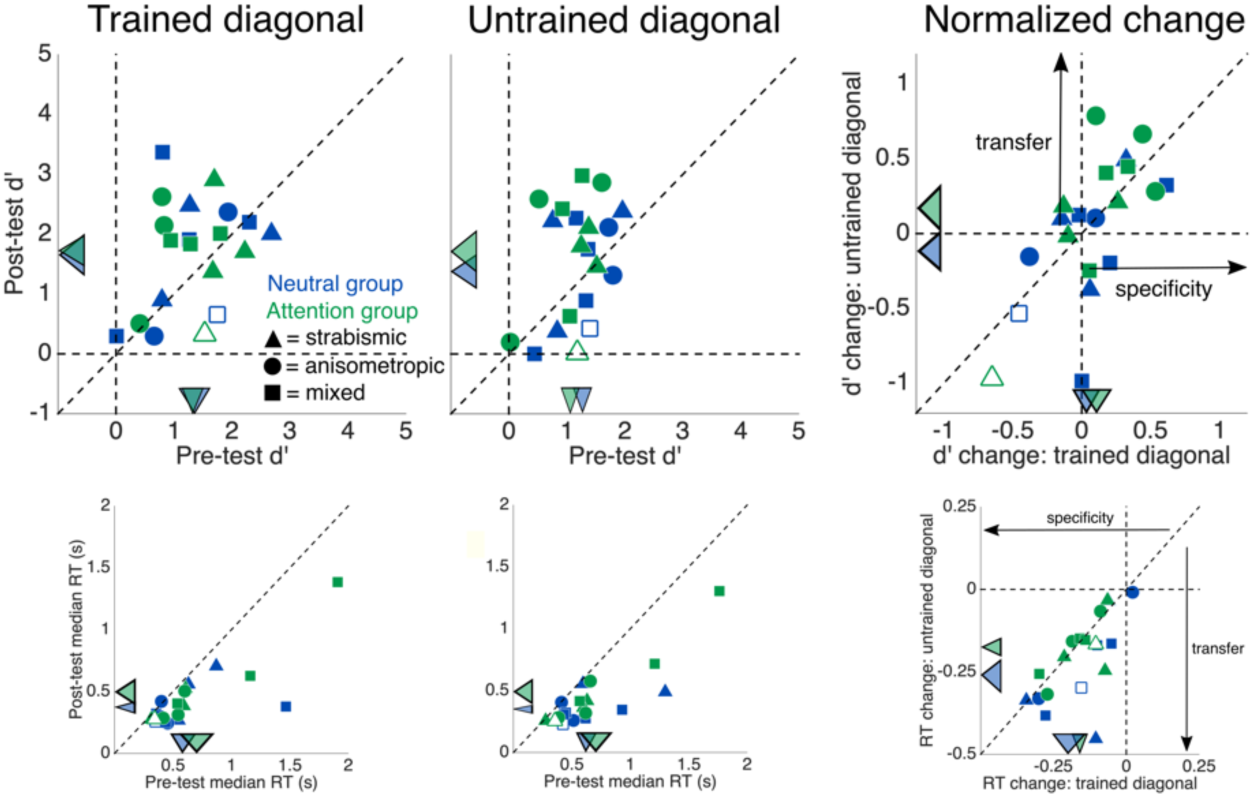
related to **Figure S1** & **S3.** Performance (top row: d’, bottom row: RT) for only the amblyopic eye of 20 individual observers on the training task at post-test versus pre-test at the trained diagonal (left column), untrained diagonal (middle column), and the relative normalized change at each diagonal (right column; data to the right of the vertical line indicate improvement at the trained diagonal; data above the horizontal line but below the unity line indicate partial transfer; data along the unity line indicate complete transfer; data above the unity line reveal greater improvement at the untrained compared to the trained diagonal). The colored arrows pointing to the axes indicate group means, and their widths represent ±1 within-groups SEM for each condition (Morey, 2008). Unfilled shapes indicate the one observer per group not included in the statistical analyses for the main training task.

**Figure S3.**
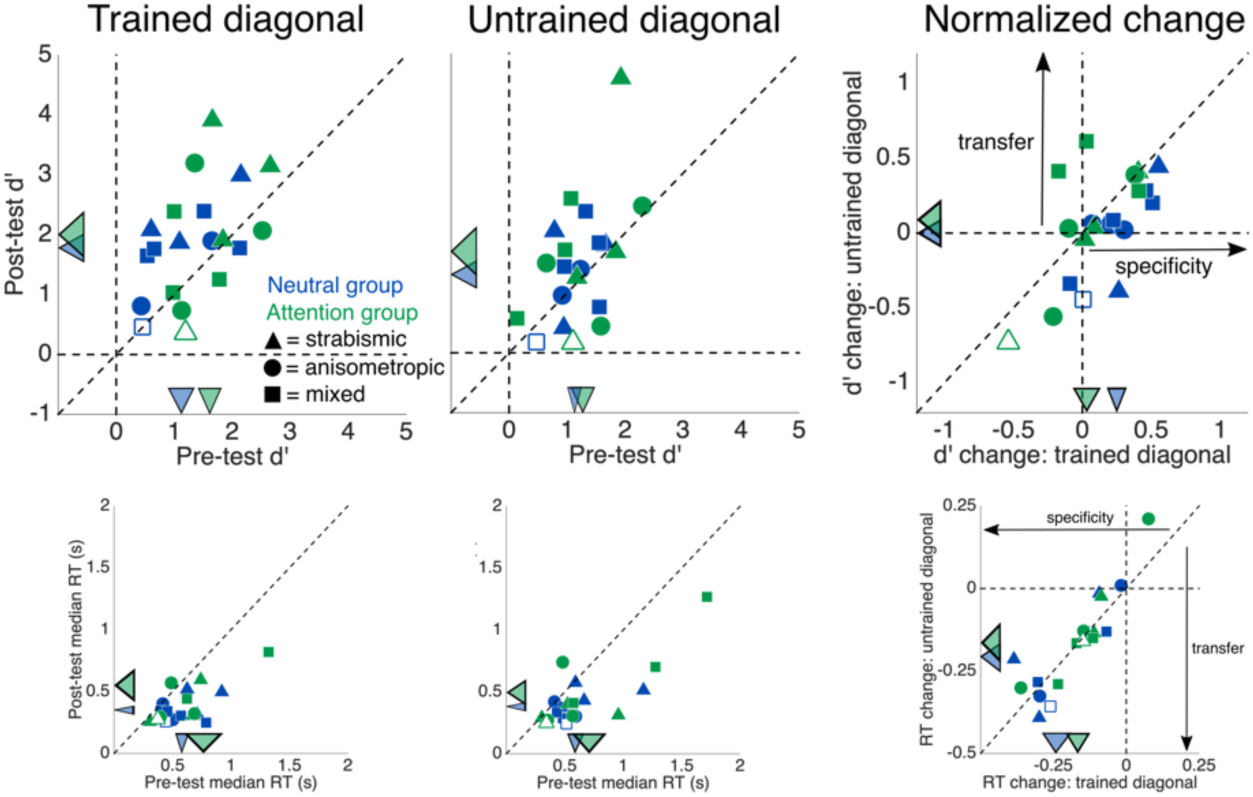
related to **Figure S1** & **S2.** Performance (top row: d’, bottom row: RT) for only the fellow eye of 20 individual observers on the training task at post-test versus pre-test at the trained diagonal (left column), untrained diagonal (middle column), and the relative normalized change at each diagonal (right column; data to the right of the vertical line indicate improvement at the trained diagonal; data above the horizontal line but below the unity line indicate partial transfer; data along the unity line indicate complete transfer; data above the unity line reveal greater improvement at the untrained compared to the trained diagonal). The colored arrows pointing to the axes indicate group means, and their widths represent ±1 within-groups SEM for each condition (Morey, 2008). Unfilled shapes indicate the one observer per group not included in the statistical analyses for the main training task.

**Figure S4.**
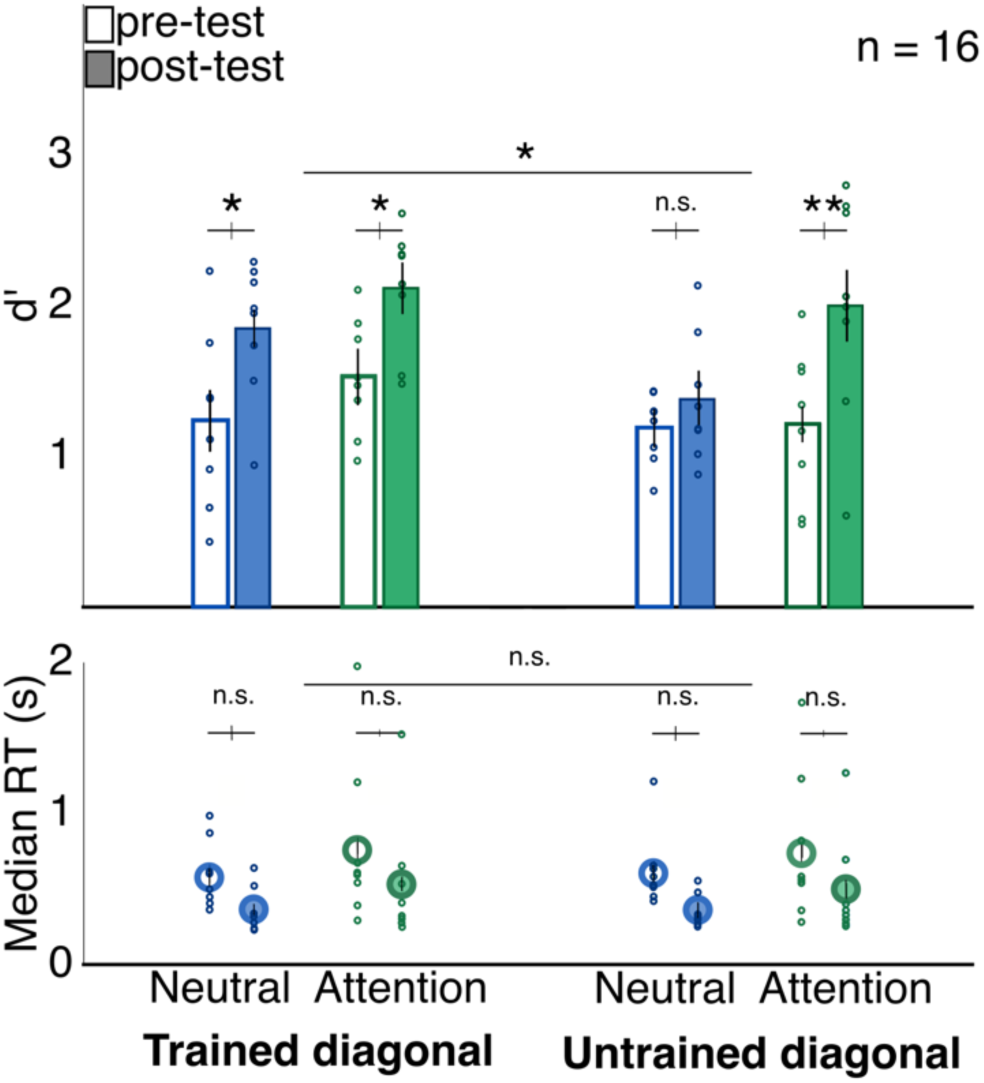
related to **Figures 3, S5**, & Table S2. Mean pre-test and post-test performance collapsed across eyes and split by diagonal, after removing the two observers who learned the least within each group. Dots indicate the performance of individual observers. Error bars represent ±1 within-subject standard error (Morey, 2008). \**p*<.05, \*\**p*<.01.

**Figure S5.**
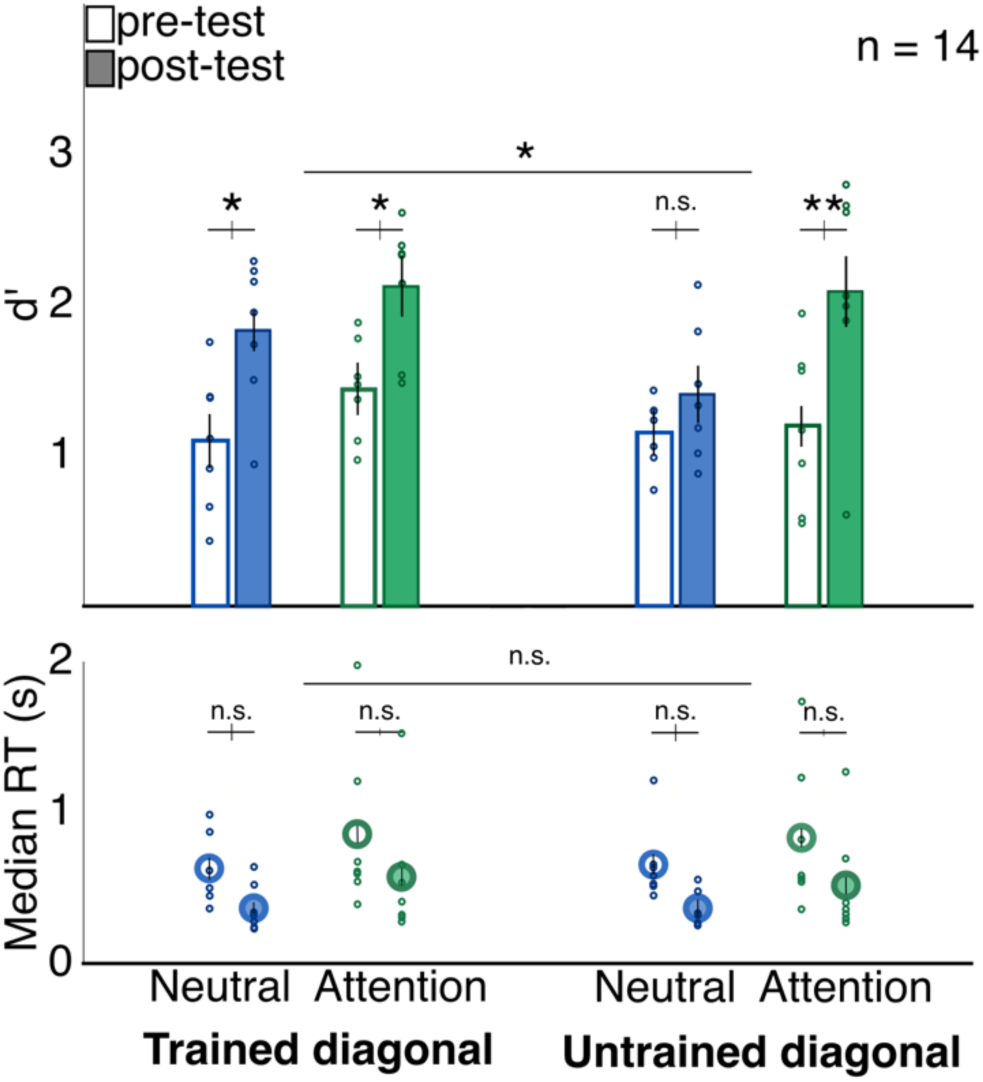
related to **Figures 3, S4** & Table S2. Mean pre-test and post-test performance collapsed across eyes and split by diagonal, after removing the three observers who learned the least within each group. Dots indicate the performance of individual observers. Error bars represent ±1 within-subject standard error (Morey, 2008). \**p*<.05, \*\**p*<.01.

**Table S2.**
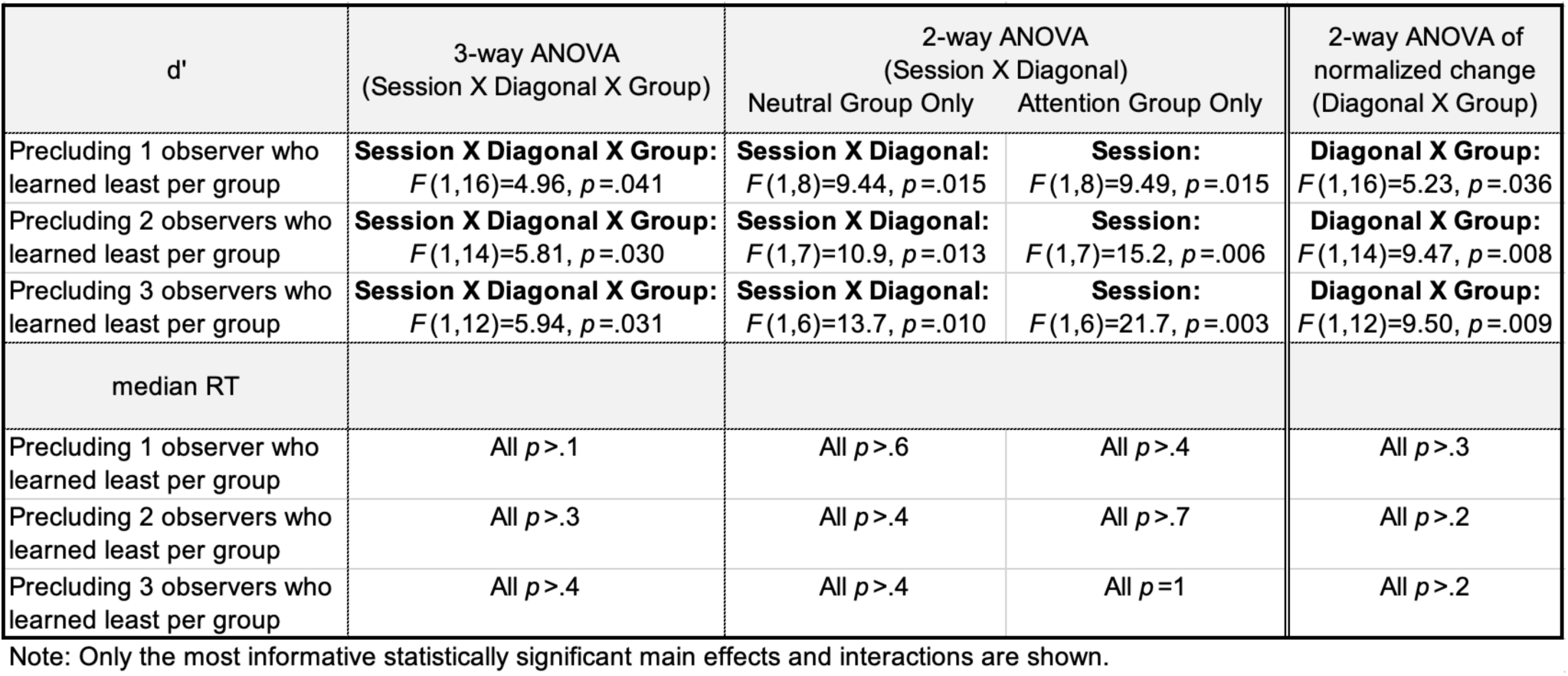
related to **Figures 3, S4** & **S5**.

| d' | 3-way ANOVA<br>(Session X Diagonal X Group) | 2-way ANOVA<br>(Session X Diagonal) |  | 2-way ANOVA of<br>normalized change<br>(Diagonal X Group) |
| --- | --- | --- | --- | --- |
|  |  | Neutral Group Only | Attention Group Only |  |
| Precluding 1 observer who<br>learned least per group | **Session X Diagonal X Group:**<br>$F(1,16)=4.96, p=.041$ | **Session X Diagonal:**<br>$F(1,8)=9.44, p=.015$ | **Session:**<br>$F(1,8)=9.49, p=.015$ | **Diagonal X Group:**<br>$F(1,16)=5.23, p=.036$ |
| Precluding 2 observers who<br>learned least per group | **Session X Diagonal X Group:**<br>$F(1,14)=5.81, p=.030$ | **Session X Diagonal:**<br>$F(1,7)=10.9, p=.013$ | **Session:**<br>$F(1,7)=15.2, p=.006$ | **Diagonal X Group:**<br>$F(1,14)=9.47, p=.008$ |
| Precluding 3 observers who<br>learned least per group | **Session X Diagonal X Group:**<br>$F(1,12)=5.94, p=.031$ | **Session X Diagonal:**<br>$F(1,6)=13.7, p=.010$ | **Session:**<br>$F(1,6)=21.7, p=.003$ | **Diagonal X Group:**<br>$F(1,12)=9.50, p=.009$ |
| median RT |  |  |  |  |
| Precluding 1 observer who<br>learned least per group | All $p>.1$ | All $p>.6$ | All $p>.4$ | All $p>.3$ |
| Precluding 2 observers who<br>learned least per group | All $p>.3$ | All $p>.4$ | All $p>.7$ | All $p>.2$ |
| Precluding 3 observers who<br>learned least per group | All $p>.4$ | All $p>.4$ | All $p=1$ | All $p>.2$ |

Note: Only the most informative statistically significant main effects and interactions are shown.

**Table S3.** related to **Figures 4 & S9**-**12**. Table of all performance values for all untrained tasks at pre-test, post-test and in terms of normalized change.

**Table S2**, related to **Figures 3, S4 & S5**.
| Neutral group | AE |  |  | FE |  |  |
| --- | --- | --- | --- | --- | --- | --- |
|  | Pre-test | Post-test | Normalized<br>change | Pre-test | Post-test | Normalized<br>change |
| Pelli Robson CS (log contrast threshold) | 1.635 | 1.845 | 0.061 | 1.890 | 1.890 | 0.001 |
| logMar acuity | 0.528 | 0.344 | 0.153 | 0.030 | 0.000 | 0.019 |
| Crowding (critical spacing in degrees of visual angle) | 0.296 | 0.322 | -0.002 | 0.107 | 0.110 | -0.014 |
| Binocular stereoacuity thresholds (arcseconds) | 899 | 738 | 0.129 | 899 | 738 | 0.129 |
| **Attention group** |  |  |  |  |  |  |
| Pelli Robson CS (log contrast threshold) | 1.767 | 1.850 | 0.026 | 1.867 | 1.900 | 0.008 |
| logMar acuity | 0.509 | 0.318 | 0.168 | 0.016 | -0.080 | 0.236 |
| Crowding (critical spacing in degrees of visual angle) | 0.423 | 0.470 | 0.002 | 0.115 | 0.116 | -0.020 |
| Stereoacuity thresholds (arcseconds) | 745 | 571 | 0.203 | 745 | 571 | 0.203 |

**Table S4.**
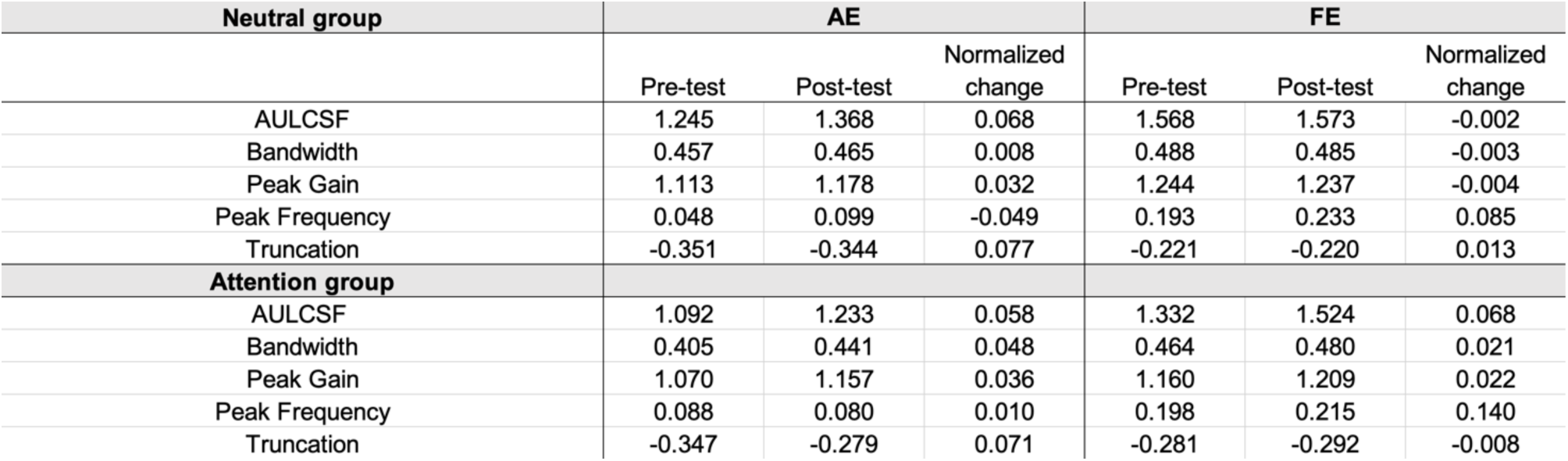
related to **Figure S7**. Table of all quick Contrast Sensitivity Function (qCSF) parameters at pre-test, post-test and in terms of normalized change.

**Figure S6.**
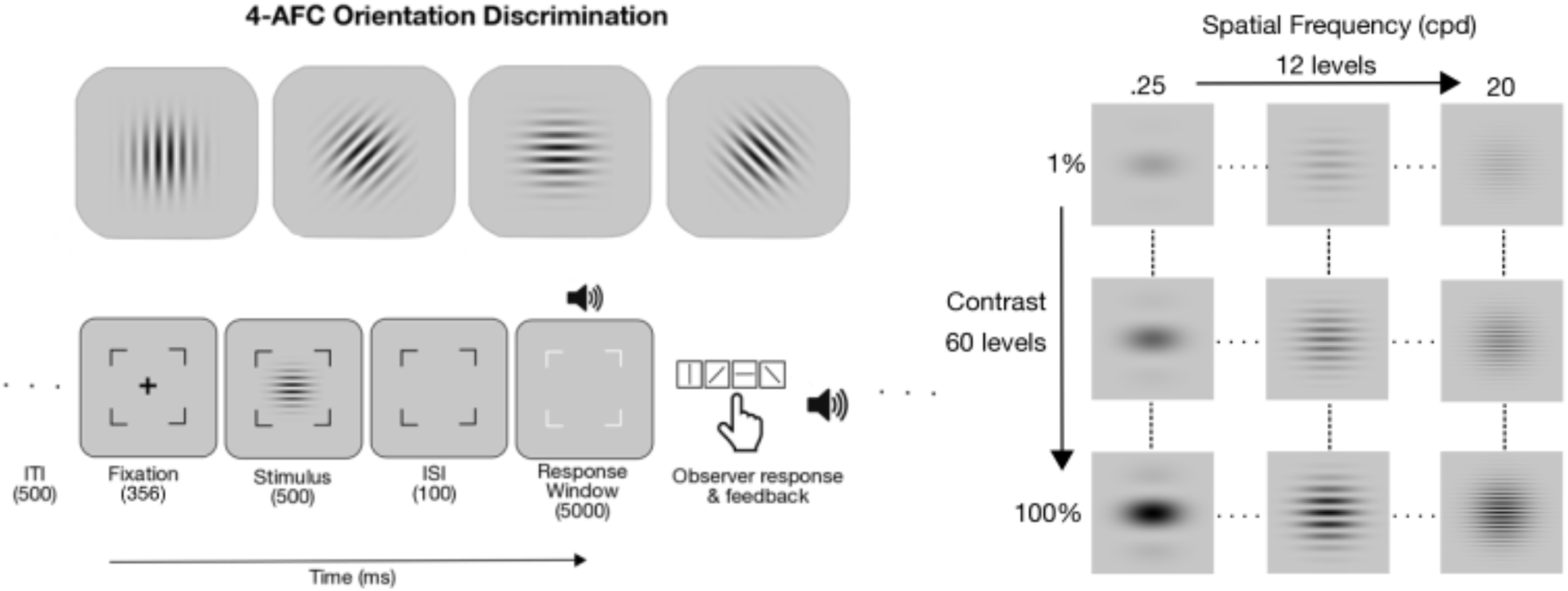
related to **STAR Methods.** The trial sequence for the qCSF behavioral task (left), and 2D array of all possible conjunctions of spatial frequency and contrast for the target stimulus (right).

**Figure S7a.**
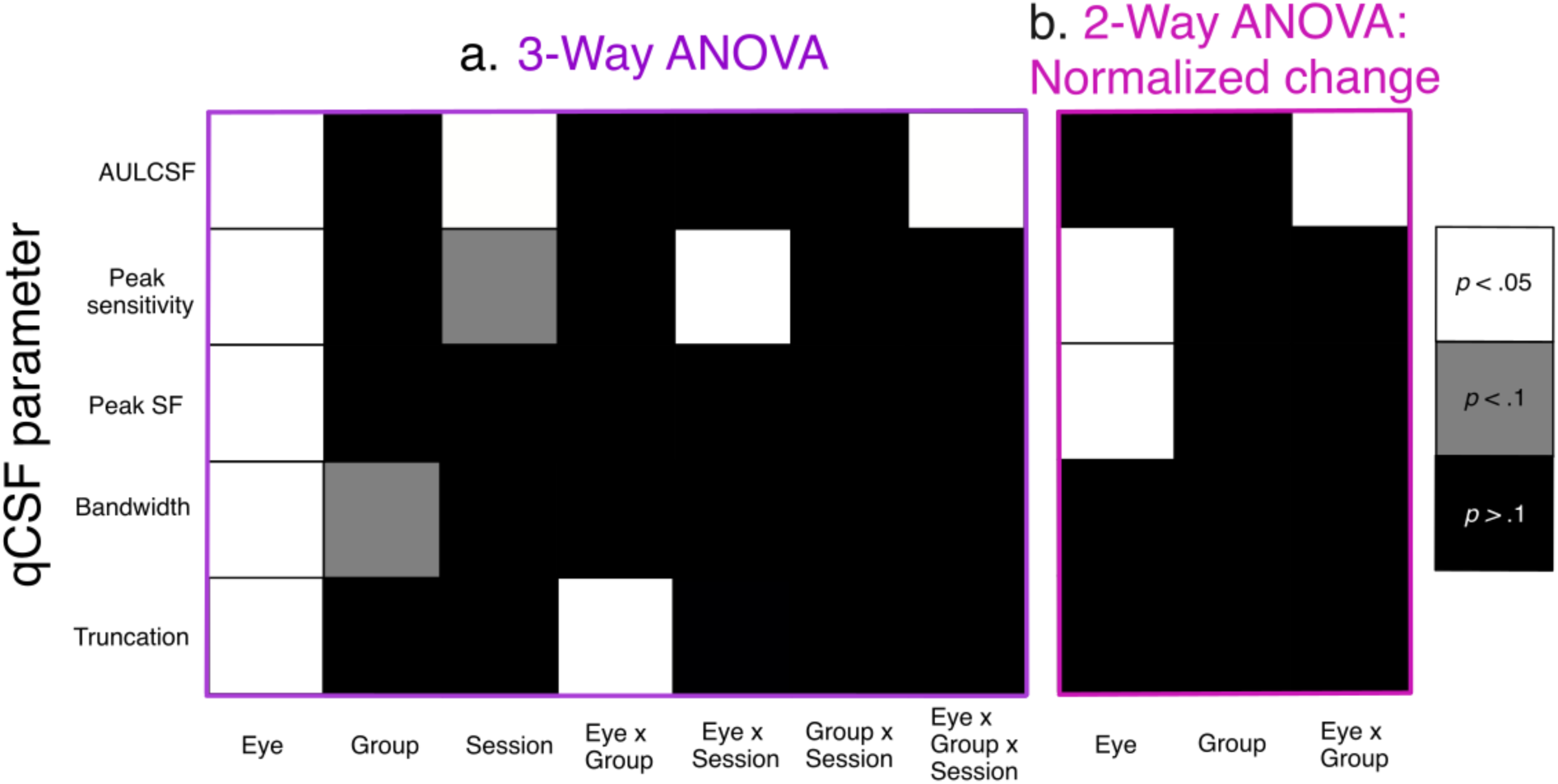
related to Table S4. A graphical representation of the results of a series of 3-way mixed ANOVAs for all qCSF parameter values (left table) and **b.** 2-WAY ANOVAs for their corresponding normalized change values (right table). The shading of each square represents the statistical significance of the main effect or interaction in each of the respective tasks.

**Figure S8.**
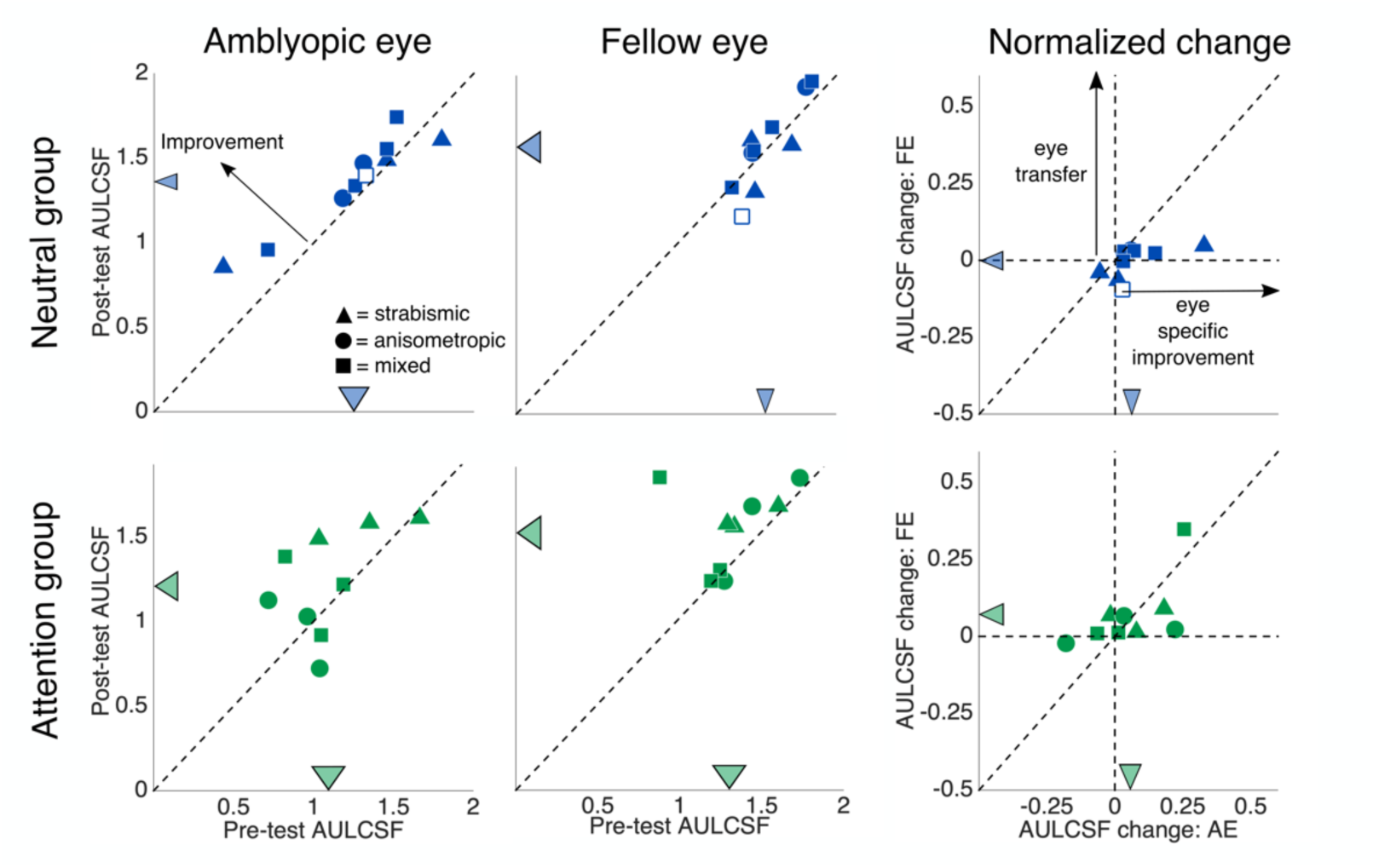
related to **Figure 4** & Table S4. AULCSF values on the qCSF task for 10 observers in the Neutral (top row) and 9 observers in the Attention group (bottom row) at post-test versus pre-test for the amblyopic eye (left column), fellow eye (middle column), and the relative normalized change in each (right column). The crossed lines symbolize group means, and their lengths indicate ±1 within-groups SEM (Morey, 2008). The unfilled shape indicates the one observer from the Neutral group not included in the statistical analyses for the main training task.

**Figure S9.**
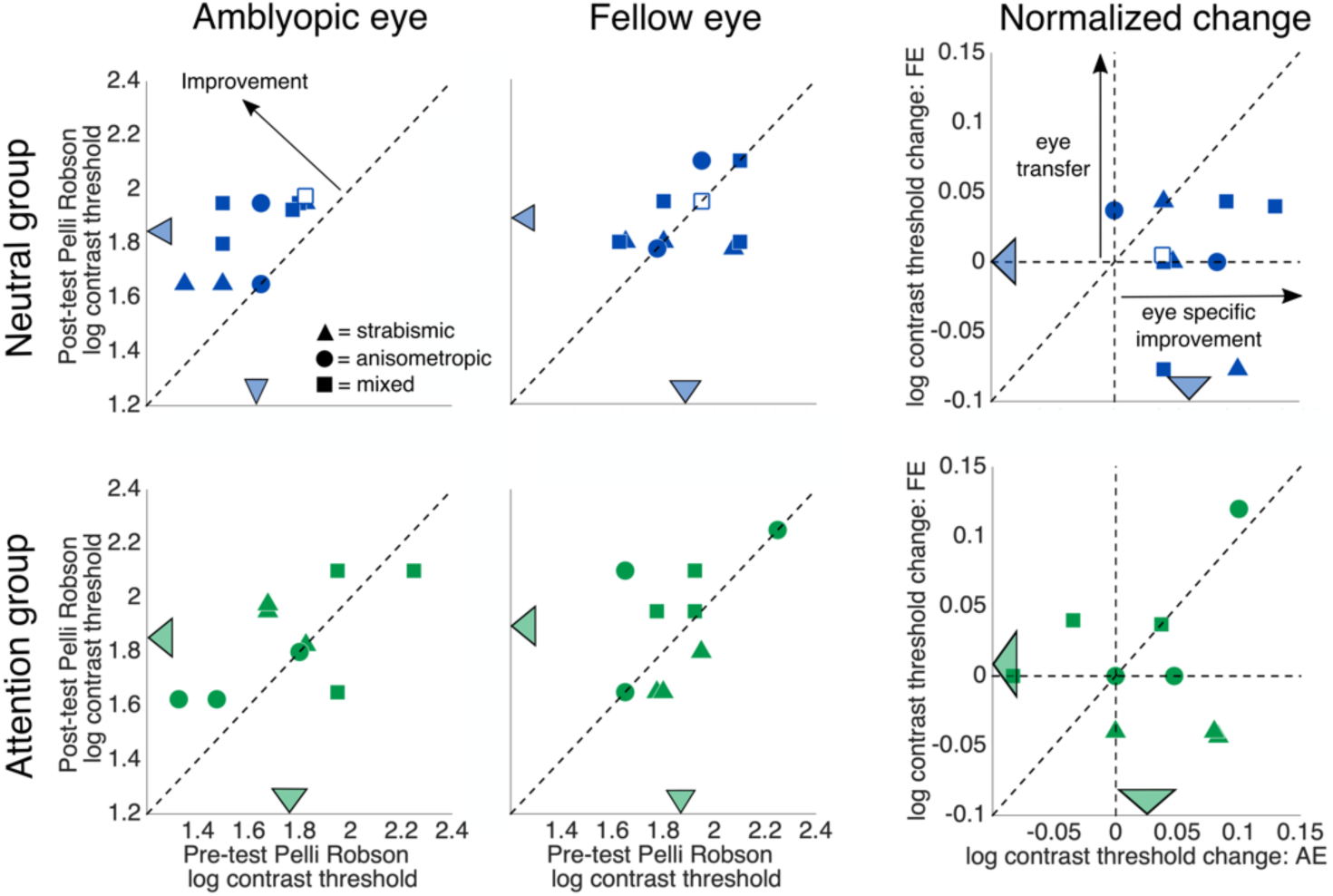
related to **Figure 4** & Table S3. Log contrast thresholds for letters on the Pelli-Robson chart for 10 observers in the Neutral (top row) and 9 observers in the Attention group (bottom row) at post-test versus pre-test for the amblyopic eye (left column), fellow eye (middle column), and the relative normalized change in each (right column). The crossed lines symbolize group means, and their lengths indicate ±1 within-groups SEM (Morey, 2008). The unfilled shape indicates the one observer from the Neutral group not included in the statistical analyses for the main training task.

**Figure S10.**
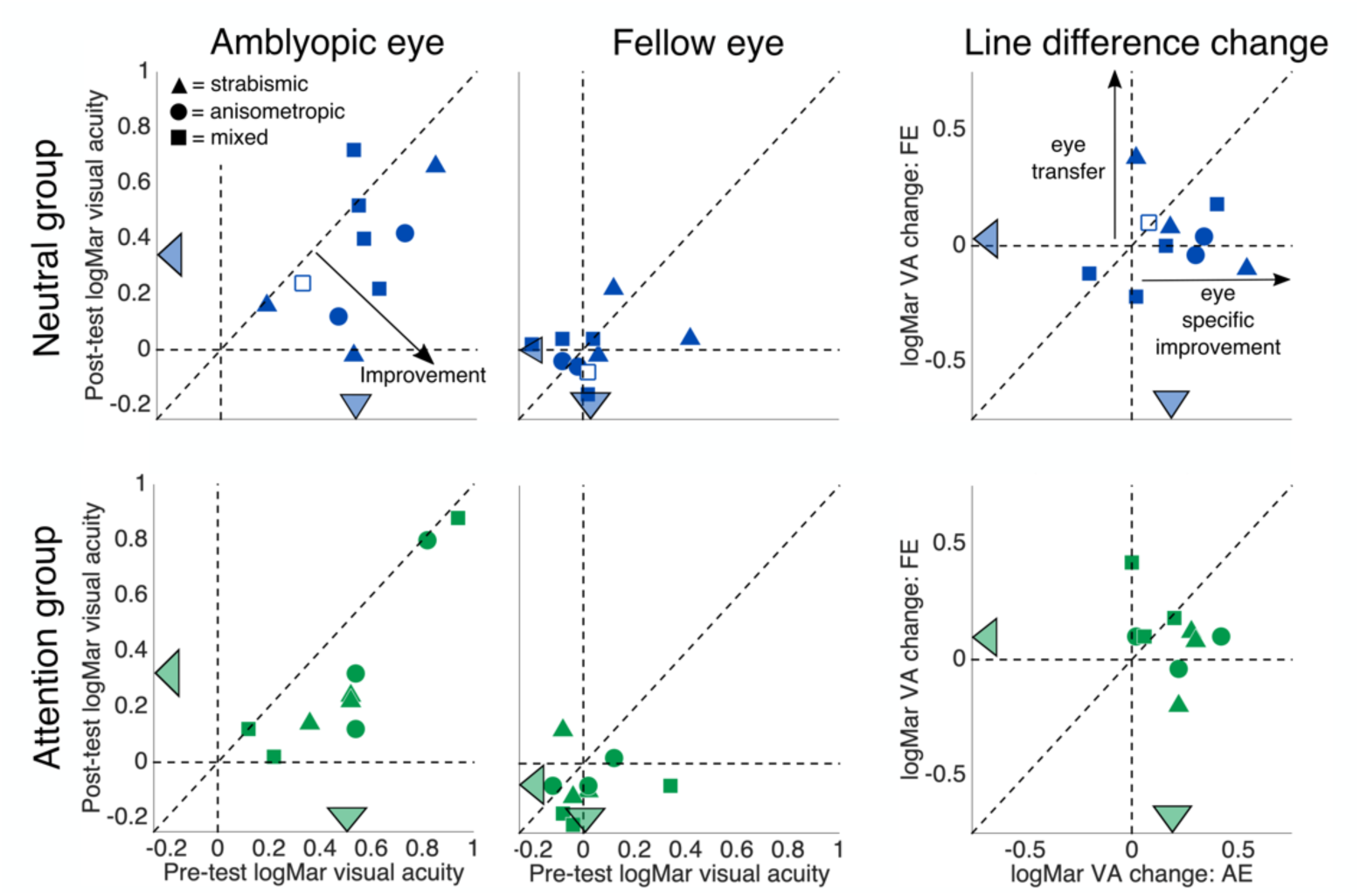
related to **Figure 4** & Table S3. logMar visual acuity for 10 observers in the Neutral (top row) and 9 observers in the Attention group (bottom row) at post-test versus pre-test for the amblyopic eye (left column), fellow eye (middle column), and the relative normalized change in each (right column). The colored arrows pointing to the axes indicate group means, and their widths represent ±1 within-groups SEM for each condition (Morey, 2008). The unfilled shape indicates the one observer from the Neutral group not included in the statistical analyses for the main training task.

**Figure S11.**
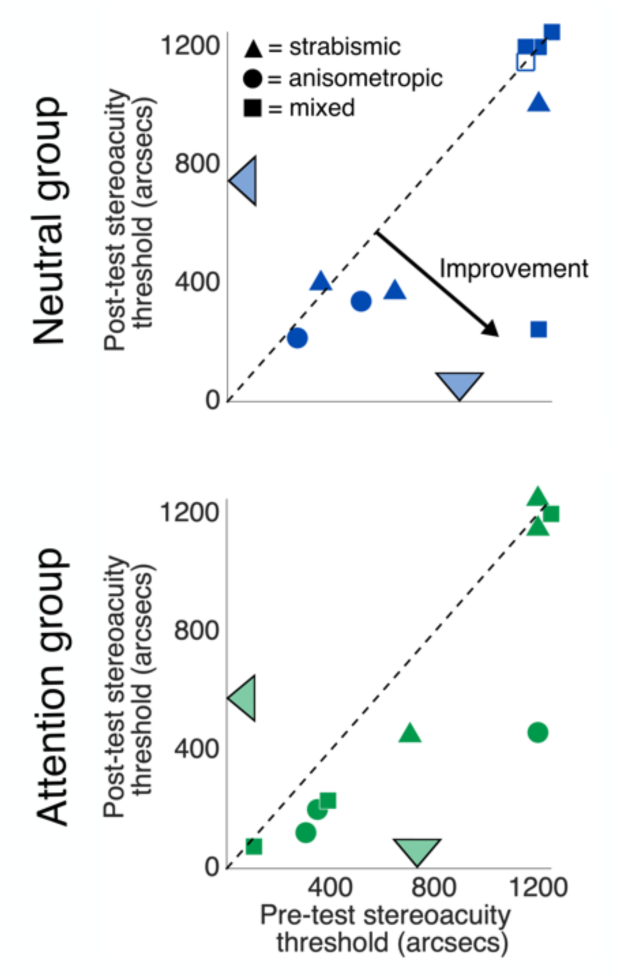
related to **Figure 4** & Table S3. Stereoacuity thresholds for 10 observers in the Neutral (top row) and 9 observers in the Attention group (bottom row) at post-test versus pre-test. The crossed lines symbolize group means, and their lengths indicate ±1 within-groups SEM (Morey, 2008). The unfilled shape indicates the one observer from the Neutral group not included in the statistical analyses for the main training task.

**Figure S12.**
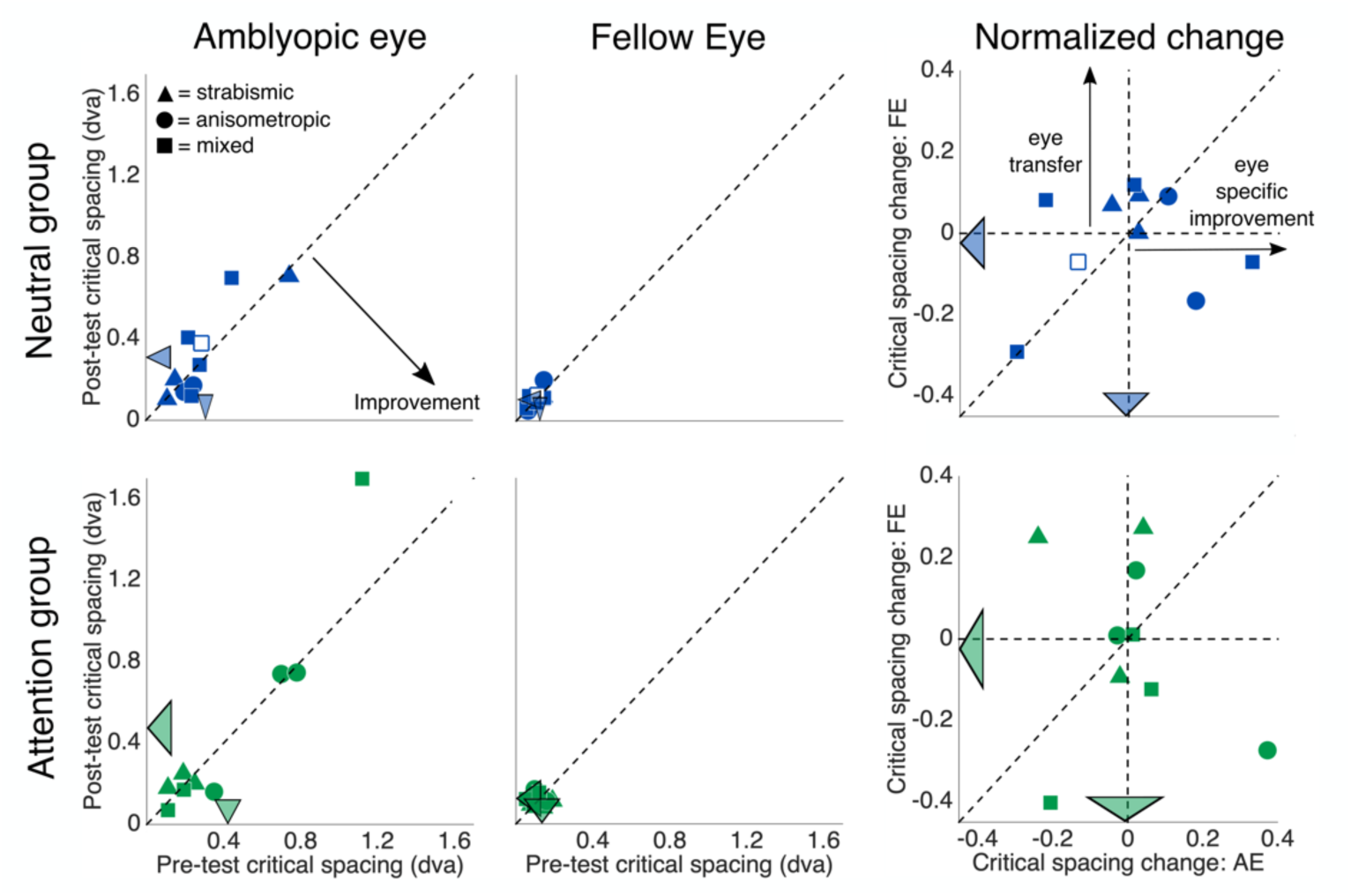
related to **Figure 4** & Table S3. Critical spacing of crowding for 10 observers in the Neutral (top row) and 9 observers in the Attention group (bottom row) at post-test versus pre-test for the amblyopic eye (left column), fellow eye (middle column), and the relative normalized change in each (right column). The crossed lines symbolize group means, and their lengths indicate ±1 within-groups SEM (Morey, 2008). The unfilled shape indicates the one observer from the Neutral group not included in the statistical analyses for the main training task.

**Figure S13.**
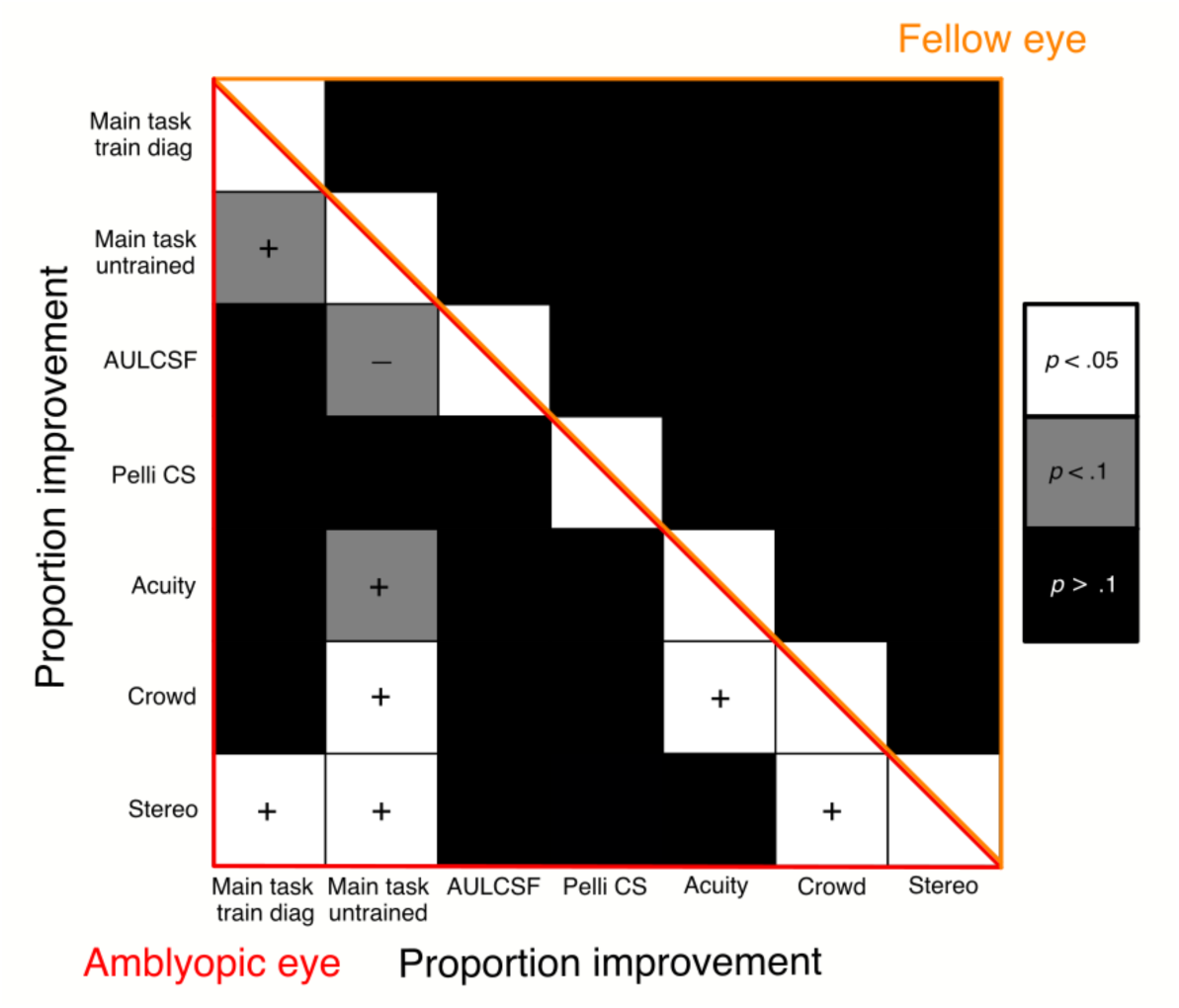
related to **Figure 2** & **4.** Correlation matrices of normalized change across all tasks after BH adjustment for the amblyopic eye (lower left triangle) and fellow eye (upper right triangle). The shading of each square represents the statistical significance of the correlation between the normalized change in each of the respective tasks. A (+) symbol indicates that greater normalized improvement in one task predicts greater normalized improvement in the other, and (-) indicates that greater improvement in one task predicts smaller improvement in the other.

## REFERENCES

Ahissar, M., Hochstein, S. (1997). Task difficulty and the specificity of perceptual learning. Nature 387, 401–406. https://doi.org/10.1038/387401a0

Ahissar, M., Nahum, M., Nelken, I., Hochstein, S. (2009). Reverse hierarchies and sensory learning. Philos. Trans. R. Soc. Lond. B. Biol. Sci. 364, 285–299. https://doi.org/10.1098/rstb.2008.0253

Asper, L., Crewther, D., Crewther, S. G. (2000). Strabismic amblyopia. Part 1. Psychophysics. Clin. Exp. Optom. 83, 49–58. https://doi.org/10.1111/j.1444-0938.2000.tb04892.x

Astle, A. T., Webb, B. S., McGraw, P. V. (2011). Can perceptual learning be used to treat amblyopia beyond the critical period of visual development? Ophthalmic Physiol. Opt. 31, 564–573. https://doi.org/10.1111/j.1475-1313.2011.00873.x

Bailey, I. L., Lovie-Kitchin, J. E. (2013). Visual acuity testing. From the laboratory to the clinic. Vision Res. 90, 2–9. https://doi.org/10.1016/j.visres.2013.05.004

Barbot, A., Landy, M. S., Carrasco, M. (2012). Differential effects of exogenous and endogenous attention on second-order texture contrast sensitivity. J Vis, 12(8), 6–6. https://doi.org/10.1167/12.8.6

Bavelier, D., Green, C. S. (2019). Enhancing attentional control: lessons from action video games. Neuron. 104, 147–163. https://doi.org/10.1016/j.neuron.2019.09.031

Benjamini Y., Hochberg Y. (1995). Controlling the false discovery rate: a practical and powerful approach to multiple testing. J. R. Stat. Soc. Series. B. Stat. Methodol. 57, 289–300. https://doi.org/10.1111/j.2517-6161.1995.tb02031.x

Brown, G. S., White, K. G. (2005). The optimal correction for estimating extreme discriminability. Behav. Res. Methods. 37, 436–449. https://doi.org/10.3758/bf03192712

Carrasco, M. (2011). Visual attention: The past 25 years. Vision Res. 51, 1484–1525. https://doi.org/10.1016/j.visres.2011.04.012

Carrasco, M. (2014). Spatial covert attention: Perceptual modulation. In A. C. Nobre & S. Kastner (Eds.), The Oxford Handbook of Attention (pp. 183–230). Oxford University Press. https://doi.org/10.1093/oxfordhb/9780199675111.013.004

Carrasco, M., Barbot, A. (2015). How attention affects spatial resolution. Cold Spring Harbor Symp. Quant. Biol. 79, 149–160. https://doi.org/10.1101/sqb.2014.79.024687

Carrasco, M., McElree, B. (2001). Covert attention accelerates the rate of visual information processing. Proc. Natl. Acad. Sci. USA 98, 5363–5367. https://doi.org/10.1073/pnas.081074098

Carrasco, M., Penpeci-Talgar, C., Eckstein, M. (2000). Spatial covert attention increases contrast sensitivity across the CSF: support for signal enhancement. Vision Res. 40, 1203–1215. https://doi.org/10.1016/S0042-6989(00)00024-9

Carrasco, M., Williams, P. E., Yeshurun, Y. (2002). Covert attention increases spatial resolution with or without masks: Support for signal enhancement. J. Vis. 2(6), 4–4. https://doi.org/10.1167/2.6.4

Cavanaugh, M. R., Zhang, R., Melnick, M. D., Das, A., Roberts, M., Tadin, D., Carrasco, M., Huxlin K. R. (2015). Visual recovery in cortical blindness is limited by high internal noise. J. Vis. 15, 9–9. https://doi.org/10.1167/15.10.9

Cavanaugh, M. R., Barbot, A., Carrasco, M., Huxlin, K. R. (2019). Feature-based attention potentiates recovery of fine direction discrimination in cortically blind patients. Neuropsychologia. 128, 315–324. https://doi.org/10.1016/j.neuropsychologia.2017.12.010

Donovan, I., Carrasco, M. (2018). Endogenous spatial attention during perceptual learning facilitates location transfer. J. Vis. 18, 7–7. https://doi.org/10.1167/18.11.7

Donovan, I., Shen, A., Tortarolo, C., Barbot, A., Carrasco, M. (2020). Exogenous attention facilitates perceptual learning in visual acuity to untrained stimulus locations and features. J. Vis. 20, 18–18. https://doi.org/10.1167/jov.20.4.18

Donovan, I., Szpiro, S., Carrasco, M. (2015). Exogenous attention facilitates location transfer of perceptual learning. J. Vis. 15, 11–11. https://doi.org/10.1167/15.10.11

Dosher, B., Lu, Z.-L. (2017). Visual perceptual learning and models. Ann. Rev. Vis. Sci. 3, 343–363. https://doi.org/10.1146/annurev-vision-102016-061249

Dugué, L., Merriam, E.P., Heeger, D.J., Carrasco, M. (2020). Differential impact of endogenous and exogenous attention on activity in human visual cortex. Sci. Rep. 10, 1–16. https://doi.org/10.1038/s41598-020-78172-x

Dugué, L., Roberts, M., Carrasco, M. (2016). Attention reorients periodically. Curr. Biol. 26, 1595–1601. https://doi.org/10.1016/j.cub.2016.04.046

Fernández, A., Carrasco, M. (2020). Extinguishing exogenous attention via transcranial magnetic stimulation. Curr. Biol. 30, 4078–4084. https://doi.org/10.1016/j.cub.2020.07.068

Gilbert, C. D., Li, W. (2012). Adult visual cortical plasticity. Neuron, 75, 250–264. https://doi.org/10.1016/j.neuron.2012.06.030

Giordano, A. M., McElree, B., Carrasco, M. (2009). On the automaticity and flexibility of covert attention: a speed-accuracy trade-off analysis. J. Vis. 9, 1–10. https://doi.org/10.1167/9.3.30

Grubb, M. A., Behrmann, M., Egan, R., Minshew, N. J., Heeger, D.J., Carrasco, M. (2013). Exogenous spatial attention: evidence for intact functioning in adults with autism spectrum disorder. J. Vis. 13, 9–9. https://doi.org/10.1167/13.14.9

Grubb, M. A., Behrmann, M., Egan, R., Minshew, N.J., Carrasco, M., Heeger, D. (2013). Endogenous spatial attention: evidence for intact functioning in adults with autism. Autism Res. 6, 108–118. https://doi.org/10.1002/aur.1269

Gu, L., Deng, S., Feng, L., Yuan, J., Chen, Z., Yan, J., Qui, X., Wang, Z., Yu, M., Chen, Z., et al. (2020). Effects of monocular perceptual learning on binocular visual processing in adolescent and adult amblyopia. iScience, 23, 100875. https://doi.org/10.1016/j.isci.2020.100875

Hariharan, S., Levi, D. M., Klein, S. A. (2005). “Crowding” in normal and amblyopic vision assessed with Gaussian and Gabor C’s. Vision Res. 45, 617–633. https://doi.org/10.1016/j.visres.2004.09.035

Harris, H., Gliksberg, M., Sagi, D. (2012). Generalized perceptual learning in the absence of sensory adaptation. Curr. Biol. 22, 1813–1817. https://doi.org/10.1016/j.cub.2012.07.059

Herrmann, K., Montaser-Kouhsari, L., Carrasco, M., Heeger, D. J. (2010). When size matters: attention affects performance by contrast or response gain. Nat. Neurosci. 13, 1554–1559. https://doi.org/10.1038/nn.2669

Hess, R. F., Thompson, B. (2015). Amblyopia and the binocular approach to its therapy. Vision Res. 114, 4–16. https://doi.org/10.1016/j.visres.2015.02.009

Hou, F., Huang, C. B., Lesmes, L., Feng, L. X., Tao, L., Zhou, Y. F., Lu, Z. L. (2010). qCSF in clinical application: efficient characterization and classification of contrast sensitivity functions in amblyopia. Investig. Ophthalmol. Vis. Sci. 51, 5365–5377. https://doi.org/10.1167/iovs.10-5468

Hou, C., Kim, Y. J., Lai, X. J., Verghese, P. (2016). Degraded attentional modulation of cortical neural populations in strabismic amblyopia. J. Vis. 16, 16–16. https://doi.org/10.1167/16.3.16

Hou F., Lesmes, L. A., Kim, W., Gu, H., Pitt, M. A., Myung, J. I., Lu, Z.L. (2016). Evaluating the performance of the quick CSF method in detecting contrast sensitivity function changes. J. Vis. 16, 18–18. https://doi.org/10.1167/16.6.18

Huang, C. B., Zhou, Y., Lu, Z. L. (2008). Broad bandwidth of perceptual learning in the visual system of adults with anisometropic amblyopia. Proc. Natl. Acad. Sci. USA 105, 4068–4073. https://doi.org/10.1073/pnas.0800824105

Hung, S. C., Seitz, A. R. (2014). Prolonged training at threshold promotes robust retinotopic specificity in perceptual learning. J. Neurosci. 34, 8423–8431. https://doi.org/10.1523/JNEUROSCI.0745-14.2014

Hussain, Z., Webb, B. S., Astle, A. T., McGraw, P. V. (2012). Perceptual learning reduces crowding in amblyopia and in the normal periphery. J. Neurosci. 32, 474–480. https://doi.org/10.1523/JNEUROSCI.3845-11.2012

Jeter, P. E., Dosher, B. A., Liu, S. H., Lu, Z. L. (2010). Specificity of perceptual learning increases with increased training. Vision Res. 50, 1928–1940. https://doi.org/10.1016/j.visres.2010.06.016

Jeter, P. E., Dosher, B. A., Petrov, A., Lu, Z. L. (2009). Task precision at transfer determines specificity of perceptual learning. J. Vis. 9, 1–13. https://doi.org/10.1167/9.3.1

Jia, W., Lan, F., Zhao, X., Lu, Z. L., Huang, C. B., Zhao, W., Li, M. (2018). The effects of monocular training on binocular functions in anisometropic amblyopia. Vision Res. 152, 74–83. https://doi.org/10.1016/j.visres.2017.02.008

Jia, K., Zamboni, E., Kemper, V., Rua, C., Goncalves, N. R., Ka Tsun Ng, A., Rodgers, C. T., Williams, G., Goebel, R., Kourtzi, Z. (2020). Recurrent processing drives perceptual plasticity. Curr. Biol. 30, 4177–4187. https://doi.org/10.1016/j.cub.2020.08.016

Jigo, M., Carrasco, M. (2020). Differential impact of exogenous and endogenous attention on the contrast sensitivity function across eccentricity. J. Vis. 20, 11–11. https://doi.org/10.1167/jov.20.6.11

Joly, O., Franko, E. (2014). Neuroimaging of amblyopia and binocular vision: a review. Front. Integr. Neurosci. 8, 62–62. https://doi.org/10.3389/fnint.2014.00062

Katz, L. M., Levi, D. M., Bedell, H. E. (1984). Central and peripheral contrast sensitivity in amblyopia with varying field size. Doc. Ophthalmol. 58, 351–373. https://doi.org/10.1007/BF00679799

Kim, S., Al-Haj, M., Fuller, S., Chen, S., Jain, U., Carrasco, M., Tannock, R. (2014). Color vision in ADHD: Part 2 - Does attention influence color perception? Behav. Brain Funct. 10, 1–9. https://doi.org/10.1186/1744-9081-10-39

Kiorpes, L., Daw, N. (2018). Cortical correlates of amblyopia. Vis. Neurosci. 35. https://doi.org/10.1017/S0952523817000232

Levi, D. M. (2006). Visual processing in amblyopia: human studies. Strabismus 14, 11–19. https://doi.org/10.1080/09273970500536243

Levi, D. M. (2013). Linking assumptions in amblyopia. Vis. Neurosci. 30, 277–287. https://doi.org/10.1017/S0952523813000023

Levi, D. M. (2020). Rethinking amblyopia 2020. Vision Res. 176, 118–129. https://doi.org/10.1016/j.visres.2020.07.014

Levi, D. M., Knill, D. C., Bavelier, D. (2015). Stereopsis and amblyopia: A mini-review. Vision Res. 114, 17–30. https://doi.org/10.1016/j.visres.2015.01.002

Levi, D. M., Li, R. W. (2009). Perceptual learning as a potential treatment for amblyopia: a mini-review. Vision Res. 49, 2535–2549. https://doi.org/10.1016/j.visres.2009.02.010

Li, H. H., Pan, J., Carrasco, M. (2021). Different computations underlie overt presaccadic and covert spatial attention. Nat. Hum. Behav., 1–14. https://doi.org/10.1038/s41562-021-01099-4

Li, R. W., Klein, S. A., Levi, D. M. (2008). Prolonged perceptual learning of positional acuity in adult amblyopia: perceptual template retuning dynamics. J. Neurosci. 28, 14223–14229. https://doi.org/10.1523/JNEUROSCI.4271-08.2008

Liu, T., Pestilli, F., Carrasco, M. (2005). Transient attention enhances perceptual performance and FMRI response in human visual cortex. Neuron 45, 469–477. https://doi.org/10.1016/j.neuron.2004.12.039

Liu, T., Stevens, S. T., Carrasco, M. (2007). Comparing the time course and efficacy of spatial and feature-based attention. Vision Res. 47, 108–113. https://doi.org/10.1016/j.visres.2006.09.017

Liu, X. Y., Zhang, T., Jia, Y. L., Wang, N. L., Yu, C. (2011). The therapeutic impact of perceptual learning on juvenile amblyopia with or without previous patching treatment. Investig. Ophthalmol. Vis. Sci., 52, 1531–1538. https://doi.org/10.1167/iovs.10-6355

Lu, Z.-L., Dosher, B. A. (2000). Spatial attention: Different mechanisms for central and peripheral temporal precues? J. Exp. Psychol. Hum. Percept. Perform. 26, 1534. https://doi.org/10.1037/0096-1523.26.5.1534

Lu, Z.-L., Li, X., Tjan, B. S., Dosher, B. A., Chu, W. (2011). Attention extracts signal in external noise: A BOLD fMRI study. J. Cogn. Neurosci. 23, 1148–1159. https://doi.org/10.1162/jocn.2010.21511

Maniglia, M., Seitz, A. R. (2018). Towards a whole brain model of Perceptual Learning. Curr. Opin. Behav. Sci. 20, 47–55. https://doi.org/10.1016/j.cobeha.2017.10.004

McKee, S. P., Levi, D. M., Movshon, J. A. (2003). The pattern of visual deficits in amblyopia. J. Vis. 3, 380–405. https://doi.org/10.1167/3.5.5

Meier, K., Giaschi, D. (2017). Unilateral amblyopia affects two eyes: fellow eye deficits in amblyopia. Investig. Ophthalmol. Vis. Sci. 58, 1779–1800. https://doi.org/10.1167/iovs.16-20964

Müller, H. J., & Rabbitt, P. M. (1989). Reflexive and voluntary orienting of visual attention: Time course of activation and resistance to interruption. J. Exp. Psychol. Hum. Percept. Perform. 15, 315–330. https://doi.org/10.1037/0096-1523.15.2.315

Morey, R. (2008). Confidence intervals from normalized data: A correction to Cousineau (2005). Tutor. Quant. Methods. Psychol. 4, 61–64. https://doi.org/10.20982/TQMP.04.2.P061

Mortazavi, M., Aigner, K. M. Antono, J. E., Gambacorta, C., Nahum, M., Levi, D. M., Föcker, J. (2020). Neural correlates of visual spatial selective attention are altered at early and late processing stages in human amblyopia. Eur. J. Neurosci. 53, 1086–1106. https://doi.org/10.1111/ejn.15024

Nakayama, K., Mackeben, M. (1989). Sustained and transient components of focal visual attention. Vision Res. 29, 1631–1647. https://doi.org/10.1016/0042-6989(89)90144-2

Nguyen, K. N., Watanabe, T., Andersen, G. J. (2020). Role of endogenous and exogenous attention in task-relevant visual perceptual learning. Plos one 15, e0237912. https://doi.org/10.1371/journal.pone.0237912

Pelli, D., Robson, J., Wilkins, A. (1988). The design of a new letter chart for measuring contrast sensitivity. Clin. Vis. Sci. 2, 187–199.

Pelli, D. G., Waugh, S. J., Martelli, M., Crutch, S. J., Primativo, S., Yong, K. X., Rhodes, M., Yee, K., Wu, X., Famira, H.F., Yiltiz, H. (2016). A clinical test for visual crowding. F1000Research 5. https://doi.org/10.12688/f1000research.7835.1

Pelli, D., Yiltiz, H. (2017). What internal noise source limits peripheral vision? J. Vis. 17, 775–775.s https://doi.org/10.1167/17.10.775

Pestilli, F., Ling, S., Carrasco, M. (2009). A population-coding model of attention’s influence on contrast response: Estimating neural effects from psychophysical data. Vision Res. 49, 1144–1153. https://doi.org/10.1016/j.visres.2008.09.018

Pham, A., Carrasco, M., Kiorpes, L. (2018). Endogenous attention improves perception in amblyopic macaques. J. Vis. 18, 11–11. https://doi.org/10.1167/18.3.11

Polat, U., Ma-Naim, T., Belkin, M., Sagi, D., (2004). Improving vision in adult amblyopia by perceptual learning. Proc. Natl. Acad. Sci. U.S.A. 101, 6692–6697. https://doi.org/10.1073/pnas.0401200101

Polat, U., Schor, C., Tong, J-. L., Zomet, A., Lev, M., Yehezkel, O., Sterkin, A., Levi, D. (2012). Training the brain to overcome the effect of aging on the human eye. Sci. Rep. 2, 278–278. https://doi.org/10.1038/srep00278

Poletti, M., Rucci, M., Carrasco, M. (2017). Selective attention within the foveola. Nat. Neurosci. 20, 1413–1419. https://doi.org/10.1038/nn.4622

Posner, M. I. (1980). Orienting of attention. Q. J. Exp. Psychol. 32, 3–25. https://doi.org/10.1080/00335558008248231

Prins, N., Kingdom. F. A. A. (2009). Palamedes: Matlab routines for analyzing psychophysical data [Computer software].

Ramesh, P. V., Steele, M. A., Kiorpes, L. (2020). Attention in visually typical and amblyopic children. J. Vis. 20, 11–11. https://doi.org/10.1167/jov.20.3.11

Roberts, M., Ashinoff, B. K., Castellanos, F. X., Carrasco, M. (2017). When attention is intact in adults with ADHD. Psychon. Bull. Rev. 25, 1423–1434. https://doi.org/10.3758/s13423-017-1407-4

Roberts, M., Cymerman, R., Smith, R.T., Kiorpes, L., Carrasco, M. (2016). Covert spatial attention is functionally intact in amblyopic human adults. J. Vis. 16, 1–19. https://doi.org/10.1167/16.15.30

Rodán, A., Marroquín, E. C., García, L. C. J. (2020). An updated review about perceptual learning as a treatment for amblyopia. J. Optom. https://doi.org/10.1016/j.optom.2020.08.002

Rolfs, M. (2009). Microsaccades: small steps on a long way. Vision Res. 49, 2415–2441. https://doi.org/10.1016/j.visres.2009.08.010

Sabesan, R., Barbot, A., Yoon, G. (2017). Enhanced neural function in highly aberrated eyes following perceptual learning with adaptive optics. Vision Res. 132, 78–84. https://doi.org/10.1016/j.visres.2016.07.011

Sagi, D. (2011). Perceptual learning in Vision Research. Vision Res. 51, 1552–1566. https://doi.org/10.1016/j.visres.2010.10.019

Schoups, A. A., Orban, G. A. (1996). Interocular transfer in perceptual learning of a pop-out discrimination task. Proc. Natl. Acad. Sci. USA 93, 7358–7362. https://doi.org/10.1073/pnas.93.14.7358

Seitz, A. R. (2017). Perceptual learning. Curr. Biol. 27, R631–R636. https://doi.org/10.1016/j.cub.2017.05.053

Sengpiel, F. (2014). Plasticity of the visual cortex and treatment of amblyopia. Curr. Biol. 24, R936–940. https://doi.org/10.1016/j.cub.2014.05.063

Sharma, V., Levi, D. M., Klein, S. A. (2000). Undercounting features and missing features: evidence for a high-level deficit in strabismic amblyopia. Nat. Neurosci. 3, 496–501. https://doi.org/10.1038/74872

Shooner, C., Hallum, L. E., Kumbhani, R. D., Ziemba, C. M., Garcia-Marin, V., Kelly, J. G., Majaj, N. J., Movshon, A., Kiorpes, L. (2015). Population representation of visual information in areas V1 and V2 of amblyopic macaques. Vision Res. 114, 56–67. https://doi.org/10.1016/j.visres.2015.01.012

Song, S., Levi, D. M., Pelli, D. G. (2014). A double dissociation of the acuity and crowding limits to letter identification, and the promise of improved visual screening. J. Vis. 14, 3–3. https://doi.org/10.1167/14.5.3

Strappini, F., Pelli, D. G., Di Pace, E., Martelli, M. (2017). Agnosic vision is like peripheral vision, which is limited by crowding. Cortex 89, 135–155. https://doi.org/10.1016/j.cortex.2017.01.012

Szpiro, S., Carrasco, M. (2015). Exogenous attention enables perceptual learning. Psychol. Sci. 26, 1854–1862. https://doi.org/10.1177/0956797615598976

Szpiro, S. F., Wright, B. A., Carrasco, M. (2014). Learning one task by interleaving practice with another task. Vision Res. 101, 118–124. https://doi.org/10.1016/j.visres.2014.06.004

Talgar, C. P., Pelli, D. G., Carrasco, M. (2004). Covert attention enhances letter identification without affecting channel tuning. J. Vis. 4(1), 3–3. https://doi.org/10.1167/4.1.3

Tsirlin, I., Colpa, L., Goltz, H. C., Wong, A. M. (2015). Behavioral training as new treatment for adult amblyopia: A meta-analysis and systematic review. Invest. Ophthalmol. Vis. Sci. 56, 4061–4075. https://doi.org/10.1167/iovs.15-16583

Vancleef, K., et al., (2019). ASTEROID: a new clinical Stereotest on an autostereo 3D tablet. Transl. Vis. Sci. Technol. 8, 25–25. https://doi.org/10.1167/tvst.8.1.25

Verghese, P., McKee, S. P., Levi, D. M. (2019). Attention deficits in amblyopia. Curr. Opin. Psychol. 29, 199–204. https://doi.org/10.1016/j.copsyc.2019.03.011

Wang, F., Chen, M., Yan, Y., Zhaoping, L., Li, W. (2015). Modulation of neuronal responses by exogenous attention in macaque primary visual cortex. J. Neurosci. 35, 13419–13429. https://doi.org/10.1523/JNEUROSCI.0527-15.2015

Wang, R., Zhang, J. Y., Klein, S. A., Levi, D. M., Yu, C. (2012). Task relevancy and demand modulate double-training enabled transfer of perceptual learning. Vision Res. 61, 33–38. https://doi.org/10.1016/j.visres.2011.07.019

Wang, R., Zhang, J. Y., Klein, S. A., Levi, D. M., Yu, C. (2014). Vernier perceptual learning transfers to completely untrained retinal locations after double training: a “piggybacking” effect. J. Vis. 14, 12–12. https://doi.org/10.1167/14.13.12

Yashar, A., Denison, R. N. (2017). Feature reliability determines specificity and transfer of perceptual learning in orientation search. PLoS Comput. Biol. 13, e1005882. https://doi.org/10.1371/journal.pcbi.1005882

Yehezkel, O., Sterkin, A., Lev, M., Levi, D. M., Polat, U. (2016). Gains following perceptual learning are closely linked to the initial visual acuity. Sci. Rep. 6, 25188. https://doi.org/10.1038/srep25188

Yotsumoto, Y., Sasaki, Y, Chan, P., Náñez Sr., J. E., Shimojo, S., Watanabe. T. (2009). Location-specific cortical activation changes during sleep after training for perceptual learning. Curr. Biol. 19, 1278–1282. https://doi.org/10.1016/j.cub.2009.06.011

Zhang, J.-Y., Cong, L.-J., Klein, S. A., Levi, D. M., Yu, C. (2014). Perceptual learning improves adult amblyopic vision through rule-based cognitive compensation. Investig. Ophthalmol. Vis. Sci. 55, 2020–2030. https://doi.org/10.1167/iovs.13-13739

Zhou, Y., Huang, C., Xu, P., Tao, L., Qiu, Z., Li, X., Lu, Z.L., (2006). Perceptual learning improves contrast sensitivity and visual acuity in adults with anisometropic amblyopia. Vision Res. 46, 739–750. https://doi.org/10.1016/j.visres.2005.07.031

